# Surface-catalyzed SAS-6 self-assembly directs centriole formation through kinetic and structural mechanisms

**DOI:** 10.1101/2020.09.04.283184

**Authors:** Niccolò Banterle, Adrian P. Nievergelt, Svenja de Buhr, Georgios N. Hatzopoulos, Charlène Brillard, Santiago Andany, Tania Hübscher, Frieda Sorgenfrei, Ulrich S. Schwarz, Frauke Gräter, Georg E. Fantner, Pierre Gönczy

**Author notes:** These authors equally contributed to this work (Niccolò Banterle, Adrian P. Nievergelt).

## Abstract

Discovering the physical principles directing organelle assembly is a fundamental pursuit in biology. Centrioles are evolutionarily conserved organelles with a 9-fold rotational symmetry of chiral microtubules imparted onto the cilia they template^1^. Centriole assemble from likewise symmetrical ring polymers of SAS-6 proteins, orthogonal to a toroidal surface surrounding the resident centriole^2–4^. How surface properties ensure ring assembly with proper symmetry and orthogonal arrangement is not known. Here, we deployed photothermally-actuated off-resonance tapping high-speed atomic force microscopy (PORT-HS-AFM) to decipher physical principles of surface-guided SAS-6 self-assembly. Using machine learning to quantify the polymerization reaction and developing a coagulation-fragmentation model, we discovered that the surface shifts the reaction equilibrium by ∼10^4^ compared to the solution situation, explaining orthogonal organelle emergence. Moreover, molecular dynamics and PORT-HS-AFM revealed that the surface converts helical SAS-6 polymers into 9-fold ring polymers with residual asymmetry, which may impart chiral features to centrioles and cilia. Overall, we discovered two fundamental physical principles directing robust centriole organelle assembly.

---

The first structure present in the nascent centriole organelle is the cartwheel, which comprises stacks of 9-fold symmetrical ring polymers ∼22 nm in diameter from which spokes emanate^2^. Each such ring polymer contains 9 homodimers of SAS-6 proteins interacting through their head domains, with a dissociation constant (K_d_) of ∼60 μM in solution^3,4^. For the human protein HsSAS-6, this is ∼10^3^ higher than the cytoplasmic concentration^5,6^, making it unclear how SAS-6 proteins can self-assemble into ring polymers in the cellular context. The nascent centriole emerges orthogonal to the surface of a torus surrounding the resident centriole to which SAS-6 is recruited (Fig. 1a, left)^7,8^. Brownian dynamics simulations suggest that such a surface might promote SAS-6 self-assembly^9^ but whether this is the case has not been addressed experimentally. To understand how weakly interacting SAS-6 homodimers can self-assemble into a ring polymer with proper symmetry, it is necessary to probe the properties of the surface polymerization reaction with high spatial and temporal resolution. Cell free assays have been quintessential for revealing the fundamental properties of other self-assembling cellular polymers, including microtubules, F-actin and FtsZ rings^10–12^. In turn, understanding fundamentals of polymer dynamics has been critical for unraveling self-organizing properties of cytoskeletal networks, and how these properties can be harnessed and modulated in vivo^13^. By analogy, unraveling SAS-6 ring polymer surface dynamics through a cell free assay is expected to shed fundamental light on the physical principles governing centriole assembly and geometry. One powerful experimental approach to probe surface protein dynamics is high-speed atomic force microscopy (AFM)^14^. We established previously that photothermally-actuated off-resonance tapping high-speed AFM (PORT-HS-AFM) is a minimally invasive method to monitor SAS-6 surface polymerization^15^. Here, we deployed PORT-HS-AFM to achieve a quantitative understanding of surface-guided SAS-6 self-assembly dynamics and thus unveil the underlying physical principles of organelle biogenesis. Purified recombinant Chlamydomonas reinhardtii SAS-6[NL] (hereafter referred to as SAS-6, Supplementary Fig. 1) formed homodimers in solution before progressively adsorbing on, and interacting with, the charged Mica surface, where PORT-HS-AFM detected the SAS-6 head domains and the spoke that emanates from each homodimer (Fig. 1b, inset). We uncovered that SAS-6 homodimers diffused on the surface and then bound to each other through their head domains, leading over time to the formation of higher order oligomers and eventually rings (Fig. 1b, Supplementary Movie 1). Initial analysis of such assembly reactions uncovered several important features. First, oligomerization occurred through incorporation of single homodimers onto pre-existing oligomers (Fig. 1c), as well as through association of higher order oligomers (Fig. 1d)^15^. Second, dissociation events also occurred for both homodimer pairs and higher order oligomers (Fig. 1e, 1f), demonstrating that the assembly reaction is reversible. Third, rings polymers containing 7 to 10 homodimers were observed, and were present in an open configuration before closing (Fig. 1i, Supplementary Fig. 2). Ring formation was also reversible, since rings could open into curved polymers without homodimer loss (Fig. 1j). Together, these findings indicate that surface-guided SAS-6 self-assembly is a complex dynamical system. Understanding how the properties of this system result in the robust formation of 9-fold symmetrical ring polymers requires a quantitative model of the self-assembly reaction. We reasoned that surface-guided SAS-6 self-assembly could be described by the coagulation-fragmentation equations, which are well suited to model reversible growth processes in physical and biological systems^16,17^. Because the SAS-6 system forms rings, such equations must be modified to include open and closed rings^9^ (Supplementary Note). The equations developed here comprise the sum of six terms: i) two terms for the appearance and disappearance of a j-mer due to coagulation (Fig. 1g), with j-mers comprising 1-10 homodimers; ii) two terms for the appearance and disappearance of a j-mer due to fragmentation (Fig. 1h); iii) two terms for ring closing and ring opening, for j-fold rings comprising 7-10 homodimers. We assumed identical association (*k*_*on*_) and dissociation (*k*_*off*_) rate constants regardless of oligomer size, as expected in a reaction-controlled system (Supplementary Note). The resulting coupled differential equations read:

**Fig. 1:**
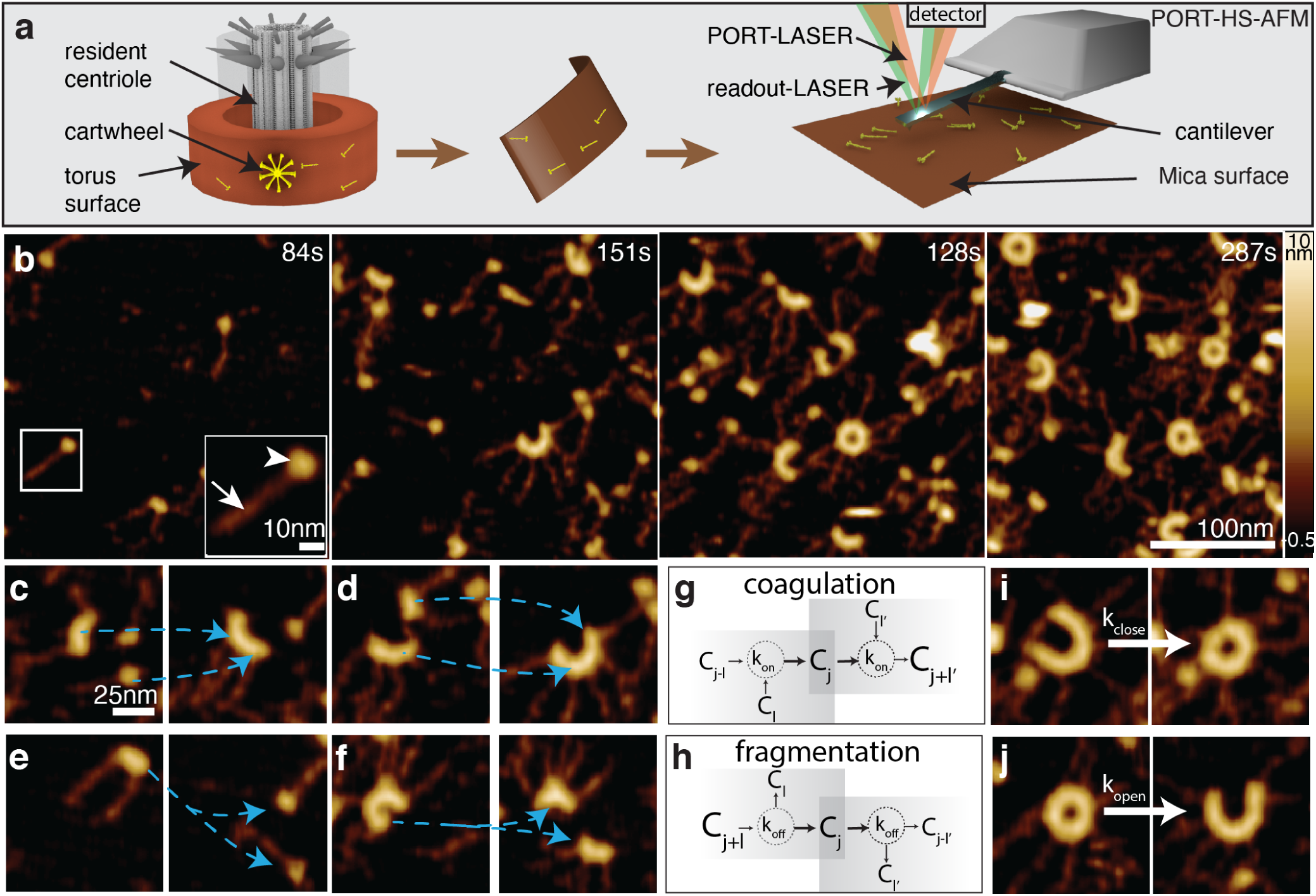
PORT-HS-AFM of surface SAS-6 self-assembly exhibits characteristics of coagulation-fragmentation system. **a**, Left; schematic of toroidal surface surrounding the resident centriole, from which the nascent centriole emerges. Middle and right: illustration of correspondence of torus surface with Mica surface in PORT-HS-AFM setting. **b**, Frames from SAS-6 self-assembly sequence monitored with PORT-HS-AFM. Higher magnification inset at 84s shows single homodimer with head domain (arrow-head) and spokes (arrow). Here and hereafter: time relative to start of recording. See also Supplementary Movie 1. **c-f**, Higher magnification of association (c, 3x+1x => 4x; d, 4x+2x => 6x) and dissociation (e, 2x => 1x+1x; f, 7 => 4x+3x) of SAS-6 homodimers monitored by PORT-HS-AFM; pairs of frames are 2.52s apart and transitions are marked by blue dashed arrows. **g, h**, Key transitions in coagulation-fragmentation system; see Equation (1) in text for details. i, j, SAS-6 ring closing (**i**) and opening (**j**) events.

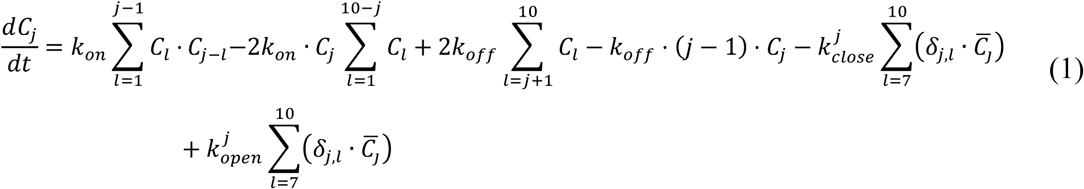

where *C*_*j*_ is the concentration of j-mers and 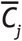 the concentration of closed j-fold rings; 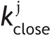 and 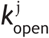 correspond to ring closing and opening rates, which potentially depend on the symmetry type j of the corresponding ring. We set out to determine parameters of this coagulation-fragmentation model experimentally. To uncover the dissociation rate *k*_*off*_, we utilized conditions with few homodimers on the surface, such that individual dissociation events were readily quantifiable, yielding 0.046 ± 0.005Hz (Fig. 2a, 2b). To determine and for each ring type, we measured the lifetime of open and closed 7- to 10-fold oligomeric species (Fig. 2c-f, Supplementary Fig. 3, Supplementary Movie 2). Interestingly, we found that the average lifetime of 9-fold_open_ was significantly shorter than that of 8-fold_open_, meaning that 9-fold_open_ polymers closed faster (Fig. 2e). Moreover, the lifetime of 7-fold_closed_ rings was shorter than that of other symmetries, accounting for their very transient nature (Fig. 2f). Together, the differences in closing and opening lifetimes lead to a kinetic stabilization of rings with 9-fold symmetry, explaining why 9-fold rings represent the majority equilibrium population^18^.

**Fig. 2:**
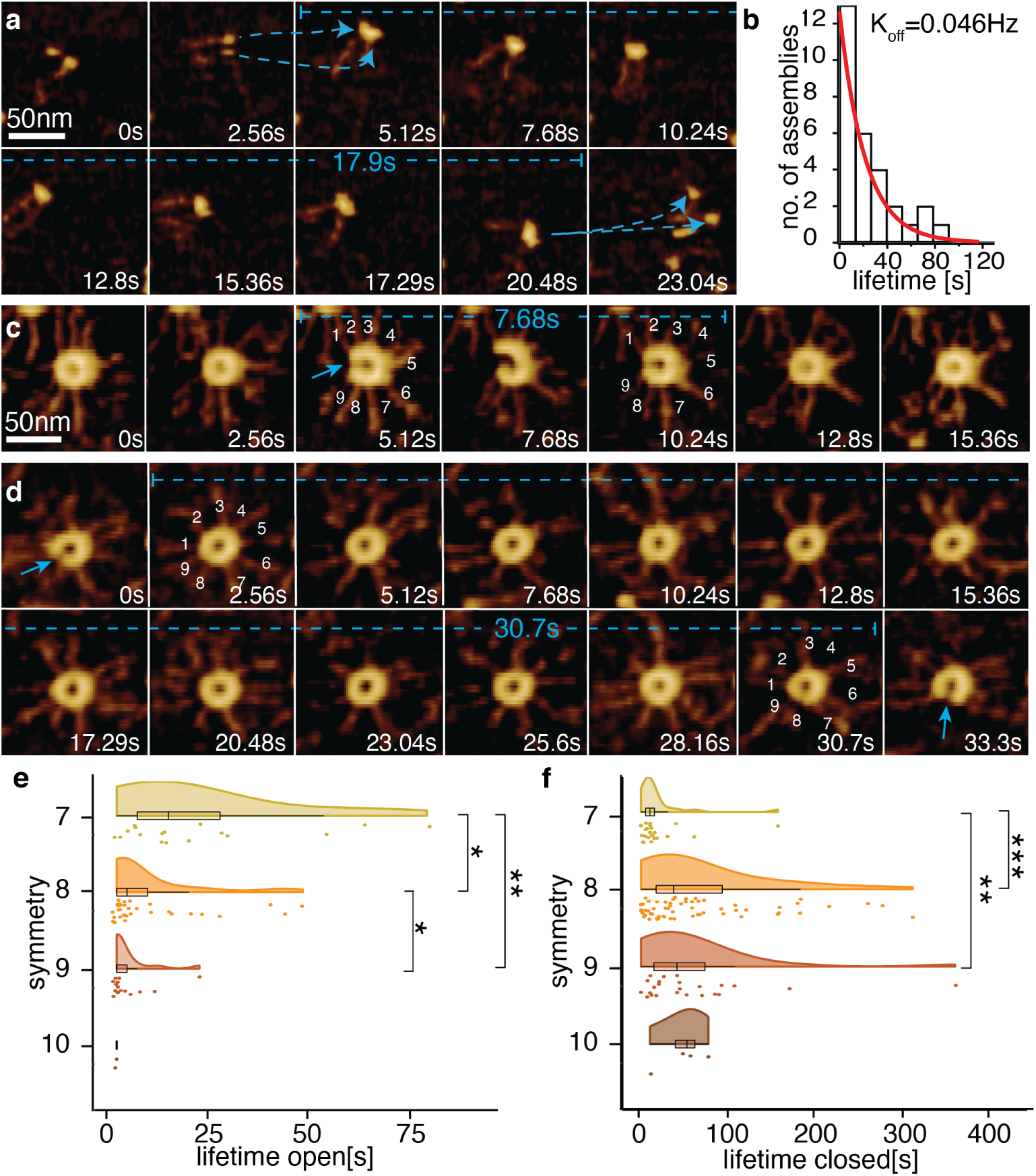
Determination of koff, 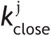 and 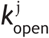 for surface SAS-6 self-assembly. **a**, Lifetime of a 2-mer analyzed with PORT-HS-AFM, forming at 5.12 s and dissociating at 20.48s (dashed blue arrows indicate association and dissociation, dashed blue line lifetime). **b**, Distribution of lifetimes of 2-mer, with exponential fitting (red line), yielding koff = 0.046 ± 0.005 Hz; n=30. **c,d**, Lifetime of open (**c**) and closed (**d**) SAS-6 rings analyzed with PORT-HS-AFM; blue arrows indicate opening events, dashed blue lines lifetimes. See also Supplementary Movie 2. **e, f**, Distribution of open (**e**) and closed (**f**) lifetimes of rings with indicated symmetries; asterisks indicate statistically significant differences (F-ratio test on the means; see Supplementary Table 1 for p-values).

Having determined 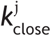 and 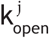, we set out to estimate *k*_*on*_ by fitting the coagulation-fragmentation equation system (1) to the experimental data, and thus compute the effective surface dissociation constant. We first determined the frequency of each oligomeric species during the assembly reaction by segmenting the PORT-HS-AFM data using machine learning aided classification (Fig. 3a, 3b, Supplementary Fig. 4, Supplementary Movie 3)^19^. We thus identified 14 classes among the 38,539 oligomeric assemblies present in five experiments and determined their prevalence over time (Fig. 3c, Supplementary Fig. 4, Supplementary Fig. 5). We then fitted these time evolutions with the coupled differential equation system (1) (Fig. 3c, Supplementary Fig. 5a-e), assessing validity with a goodness of fit test and optimality versus both simpler and more complex models (Supplementary Fig. 6). The fitting yielded an effective surface dissociation *K*_*d*_*= k*_*off*_*/k*_*on*_ constant of 79 ± 12 homodimers/μm^2^. In human cells, ∼100 HsSAS-6 homodimers are present at centrioles^5^, whereas the surface of the torus is ∼0.1μm^2 (20)^, translating into an effective surface concentration of ∼1000 homodimers/μm^2^, ten times higher than the *K*_*d*_ determined here. Therefore, the catalytic effect conferred by the surface shifts the equilibrium by a factor ∼10^4^ compared to the solution situation, where the concentration is ∼10^3^ lower than the K_d_. This provides an explanation for how rings can assemble exclusively on the surface of the torus and not in the cytosol. Overall, we conclude that surface-guided catalysis surmounts the weak interaction and low concentration that homodimers exhibit in solution, thus ensuring efficient SAS-6 self-assembly.

**Fig. 3:**
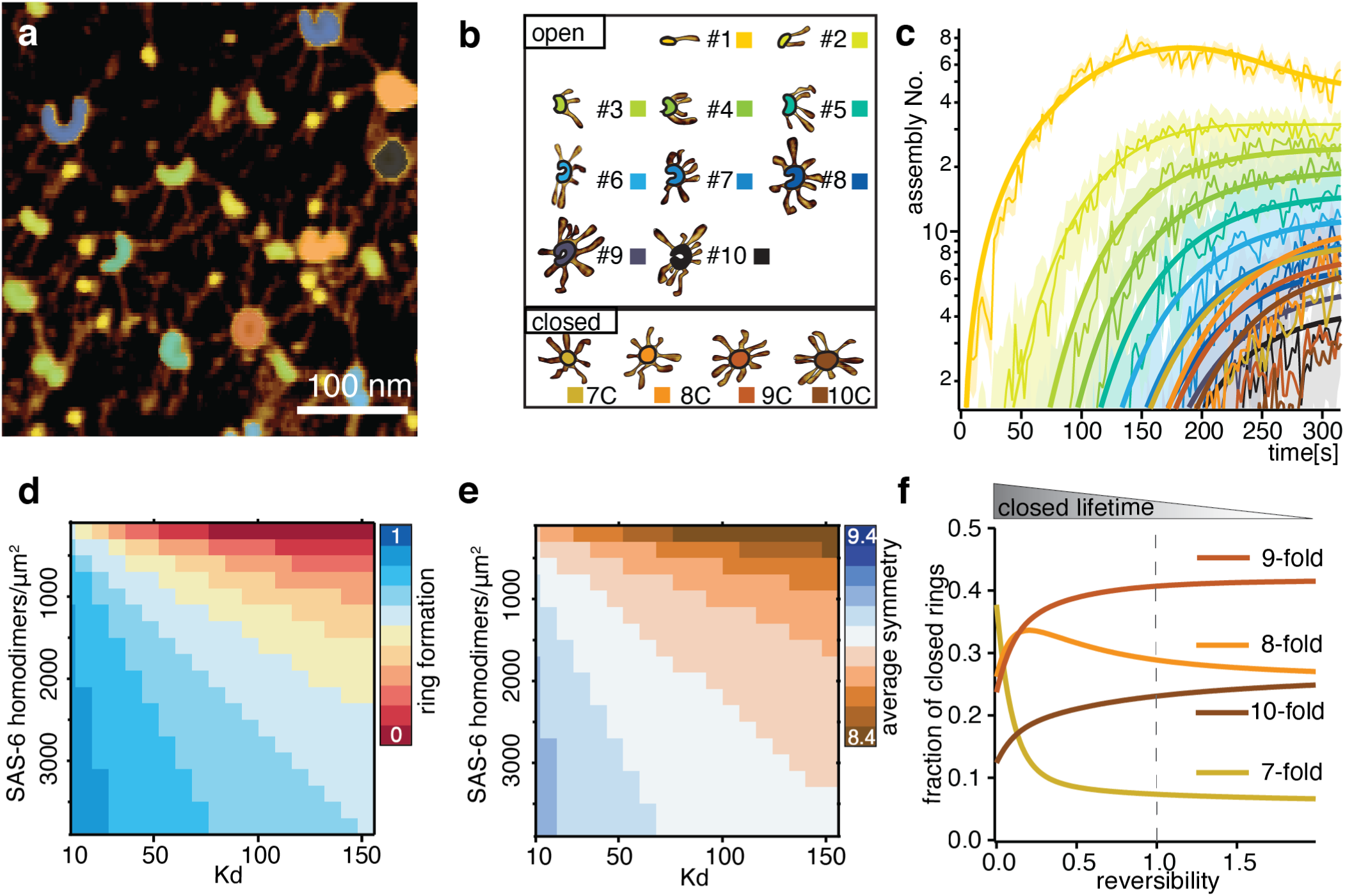
Machine learning aided classification and fitting of surface SAS-6 self-assembly with coagulation-fragmentation model. **a**, Distinct oligomeric states recognized by machine learning aided classification in PORT-HS-AFM data. **b**, Examples of the existing 14 classes, some of which are present in panel a. See also Supplementary Movie 3. **c**, Time evolution of each oligomeric species (thin lines, colors as in panel B) in one PORT-HS-AFM experiment fitted with coagulation-fragmentation model (overlaid thick lines). d, e, Model predictions of ring formation efficiency (**d**) and average ring symmetry (**e**) at varying Kd and SAS-6 homodimers/μm2. f, Fraction of closed rings of indicated symmetries calculated as numerical solution of the fitted model after 17 min as a function of closed lifetime (0: irreversible closing, 1: actual experimental values).

We used the above quantitative model of surface-guided SAS-6 self-assembly to computationally explore the impact on ring formation efficiency and on ring symmetry of changing parameters that could be modulated in the cellular context. We reasoned that the effective surface concentration of SAS-6, the effective dissociation constant between homodimers, as well as the lifetimes of ring opening and closing could each be such modulation targets. Varying these parameters in a computational analysis of our physical model, we uncovered that ring formation efficiency exhibited a strong dependence on effective surface concentration and dissociation constant (Fig. 3d). This finding can explain why supernumerary centrioles form in human cells expressing HsSAS-6 in excess^21^, which likely increases the effective surface concentration on the torus^6^. We found by contrast that ring symmetry at equilibrium was remarkably robust to parameter changes, remaining close to 9-fold within the entire explored range (Fig. 3e). Symmetries also remained essentially unchanged when closed lifetimes became shorter, which mimics enhanced reversibility (Fig. 3f, right-most). Interestingly, however, when the lifetime of closed rings became longer, which mimics poor reversibility, symmetry shifted towards 7-fold due to kinetic trapping (Fig. 3f, left-most). This result stresses the necessity of reversible association and dissociation to avoid such kinetic trapping^22^, raising the intriguing possibility that this may explain why a weak interaction between SAS-6 head domains has been maintained across evolution. Overall, we conclude that symmetry-dependent open and closed lifetimes, as well as reversibility of the assembly reaction, together ensure proper ring symmetry over a large parameter range.

What are the physical reasons for symmetry-dependent open and closed lifetimes? The fundamental parameters dictating opening and closing of any ring are the native curvature and flexibility of its constituent polymer^23^. To evaluate these parameters in the case of SAS-6, we analyzed intermediate assembly states, where curvature is not influenced by closing constraints, because reactive ends are far from one another (Supplementary Fig. 7). We found that the angle between neighboring SAS-6 homodimers in 4- and 6-mers reflects a curvature that would yield rings with a symmetry between 9- and 10-fold, with polymer flexibility reflected by the standard deviation of the curvature (Fig. 4a). To explore how such polymer characteristics could result in a closed ring, we turned to the crystal structure of the SAS-6 homodimer with 6 heptad repeats of the coiled-coil (hereafter referred to as SAS-6[6HR]). Interestingly, we found that whereas computational assembly of 9 such homodimers yielded a flat ring when enforced by modeling (Fig. 4b, blue)^3^, without imposing constraints it yielded instead a shallow spiral with a ∼7.9 nm pitch (Fig. 4b, red). Such a spiral is reminiscent of the helical filaments formed by the structurally related proteins CCDC61 and XRCC4^24^, as well as of the steep SAS-6 spiral proposed in C. elegans centrioles^25^. How could a molecule that might form an intrinsically helical, and thus chiral, structure assemble into a ring on a surface? A theory of the preferred conformation of a helix when constrained on a surface predicts the existence of metastable states, including a signature S shaped conformation^26,27^. Remarkably, PORT-HS-AFM indeed uncovered instances of such transient conformations for SAS-6 (Fig. 4c-f, Supplementary Movie 1). Taken together, these findings suggest that SAS-6 polymers are intrinsically helical but constrained into ring polymers upon surface interaction. To gain quantitative insights into how an intrinsically helical SAS-6 polymer forms a ring on a surface, we performed extensive coarse-grained molecular dynamics (MD) computer simulations of SAS-6[6HR] in different oligomerization states, both in solution and on a surface. The resulting distribution of in-plane angles between homodimers in solution revealed that the conformational ensemble for 4- and 6-mers would result in a ∼6-fold symmetry (Fig. 4g, Supplementary Fig. 8). Importantly, we found that the surface significantly lowered such in-plane angles, bringing them close to a configuration enabling 9-fold symmetry (Fig. 4g, Supplementary Fig. 8), in agreement with the experimental value (Fig. 4a). Therefore, the surface plays a critical role in promoting in-plane angles close to 40° between neighboring SAS-6 homodimers, thus favoring 9-fold symmetrical ring polymers. Remarkably, another feature unveiled by the MD simulations is that SAS-6[6HR] rings were not entirely flat, since spokes were either bound or not bound to the surface (Fig. 4h), as evident from the torsional angle between homodimers (Fig. 4i, Supplementary Fig. 8). Furthermore, the spatial distribution of spokes that are either bound or not bound was not random (Fig. 4j, p<0.0001); instead, the behavior of neighboring spokes tended to be correlated, likely because the initial helical nature of the SAS-6 polymer imparts asymmetry to the ring. Analyzing spoke mobility as a proxy for surface interaction of 9-fold SAS-6 rings in the PORT-HS-AFM data set supported this conclusion (Supplementary Fig. 9). Overall, we conclude that the presence of a surface imparts an inherent asymmetry to the SAS-6 ring polymer. In conclusion, our findings reveal core biophysical principles fundamental for the onset of centriole biogenesis. We demonstrated that surface guided SAS-6 self-assembly is a coagulation-fragmentation reaction in which all intermediates react with one another, with dynamic ring opening and closing events being critical for robust generation of 9-fold symmetrical rings. Furthermore, we unveiled that the surface catalyzes SAS-6 ring assembly, shifting the equilibrium by a factor of ∼10^4^ from that in solution. This finding reinforces the emerging understanding that the interplay between solution and surface compartments, for instance between cytosol and membrane, characterizes many cellular diffusion-reaction systems^28^, and reveals that this principle is relevant at the space scale of organelle assembly. From a geometrical point of view, the surface catalysis uncovered here is reminiscent of that acting at membranes to promote the formation of macromolecular complexes such as pore forming toxins^29,30^. In the cellular context, the torus surface is thus expected to efficiently restrict SAS-6 ring polymer assembly to the vicinity of the resident centriole. Given that this surface is parallel to the long axis of the resident centriole (see Fig. 1a), this could explain how orthogonal emergence of the nascent centriole is achieved. Furthermore, given that surface-constrained SAS-6 ring polymers exhibit spoke asymmetry, we speculate that this might impart the universal signature chirality of centriolar and ciliary microtubules.

**Fig. 4:**
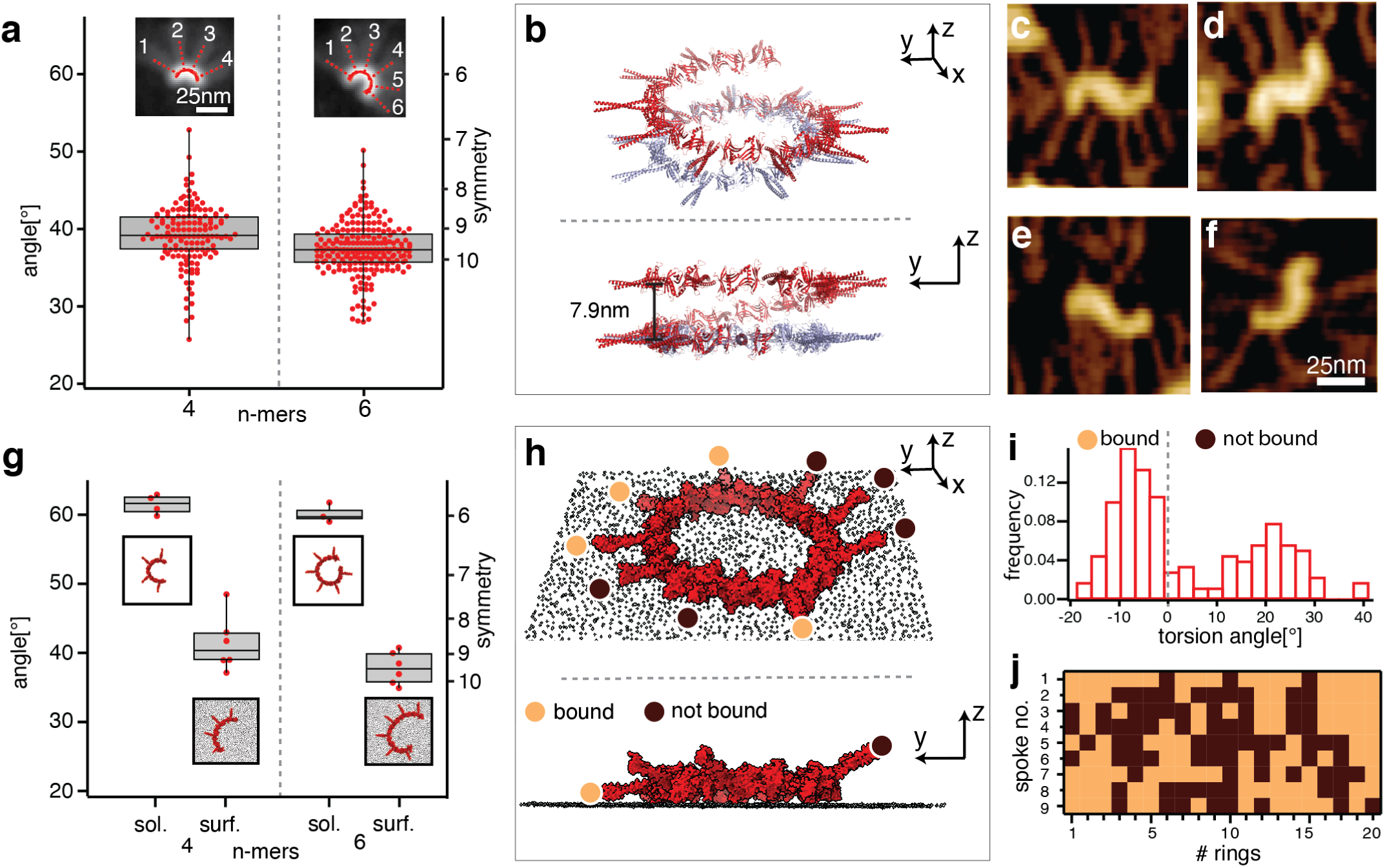
SAS-6 ring polymers on the surface exhibit asymmetric features. **a**, Box plot angle distribution of SAS-6 4- and 6-mers from PORT-HS-AFM data set (mean ± SD: α_4_ = 39°± 4°, α_6_ = 38°± 3°). Insets: averaged and realigned images with fit (solid red line, dashed red lines denote spoke locations). **b**, Isometric (top) and side (bottom) views of 9-mer generated from SAS-6[6HR] crystal structures, resulting in a spiral (red); the closed ring (blue) is enforced by modeling5. **c-f**, Examples of transient S-shaped SAS-6 oligomers observed with PORT-HS-AFM. **g**, Box plots angle distribution from MD simulations of SAS-6[HR] 4- and 6-mers in solution and on a surface. Insets show examples of corresponding simulated structures. **h**, Isometric (top) and side (bottom) view from MD simulation of closed SAS-6[HR] 9-mer on a surface, with spokes either bound to the surface (beige disks) or not bound (dark brown disks); only two spokes are highlighted in the side view. **i, j**, Torsion angle distribution in 20 closed SAS-6[HR] 9-mers on a surface simulated by MD (**i**), and corresponding matrix representation (**j**), with spokes classified as bound or not bound (bound when torsion angle <0, dashed line in i).

## Supporting information

Supplementary Movie 1: SAS-6 oligomerization and ring assembly

Supplementary Movie 2: SAS-6 ring opening and closing

Supplementary Movie 3: Classification result from machine learning software Ilastik

## Methods

### Protein expression and purification

His-tagged CrSAS-6[NL] spanning amino acids 1-503 of the protein (see Supplementary Fig. 1a)^1^ was expressed in the *Escherichia coli* strain BL21(DE3) (Stratagene). Bacteria were grown at 37 °C in lysogeny broth (LB) supplemented with kanamycin (50μg/mL) up to an absorption *A*_600_ of 0.7, when 0.3 mM IPTG (isopropyl β-D-1-thiogalactopyranoside) was added to induce protein expression overnight at 18 °C with shaking at 210 rpm. Cells were collected by centrifuging for 10 min at 4 °C at 4,000*g* (JLA-9.1000, Beckman Coulter). The bacterial pellet was resuspended in 50 mM Tris-HCl pH 8.0, 400 mM NaCl, 20 mM Imidazole pH 8.0, 3 mM β-mercaptoethanol (β-ME), 50 Units DNAseA, 3mM MgCl2, protease inhibitors (Complete EDTA-free, Roche) and 1% v/v Tween 20, and lysed by sonication (100 cycles of 3 s pulses followed by 12 s breaks, with the sample kept in ice). Cellular debris were removed by centrifugation at 49,000*g* (JA 25.50. Beckman Coulter) for 1 h at 4 °C. For nickel purification, the cleared lysate was loaded at 4 °C on an HisTrap HP Ni^2+^–Sepharose column (GE Healthcare) following the manufacturer’s instructions and further purified by size exclusion chromatography (SEC) using a HiLoad Superdex 200 16/60 column (GE Healthcare) equilibrated in 20 mM Tris-HCl pH 7.4, 150 mM KCl. PreScission protease was added to remove the tag followed by an additional reverse HisTrap step to remove un-cleaved protein prior to SEC. After verifying purity with Coomassie stained SDS-PAGE (see Supplementary Fig. 1b), the sample was concentrated to 1-3mg/mL, depending on the specific preparation, with an Amicon ultra filter unit with a 10 kDa cut-off, snap-frozen in liquid nitrogen into aliquots and stored at –20 °C (*1*)

### PORT-HS-AFM imaging

PORT-HS-AFM imaging was performed on a custom-built high-speed setup with a head scanner and controller assembled on a spring-based vibration isolator (BM4, Minus-K), around a commercial Multimode 8 AFM base, combined with a piezo amplifier (Techproject), as previously described^2^. The entire setup was kept at 6–10 °C by placing the microscope base, scanner and head into a low-vibration cooler (WL450F-220-FL, Swisscave). Cantilever tips were electron-beam deposited from a carbon precursor (Tetradecane, Sigma-Aldrich 172456) onto the existing silicon nitride tips (BL-AC10DS, Olympus) to reduce protein adhesion to the cantilever and allow their recycling. For imaging, 60 µl of imaging buffer (20 mM Tris-HCl, 150 mM KCl, pH 7.4) were injected into the cantilever holder of the head. After aligning the lasers, the head was placed onto the scanner directly after cleaving the glued-on 3 mm Muscovite Mica disk (Electron Microscopy Sciences, Hatfield). The concentrated sample was diluted in imaging buffer to 60 µg/mL. After pre-loading the Hamilton syringe with 15 µl of imaging buffer to compensate for the dead volume, 5 µl of the diluted sample was directly injected in the liquid cell already containing 60 µL of buffer, reaching a final concentration of 4 µg/mL (∼60nM). For measurement of ring opening and closing lifetimes, a 10 µL droplet of sample diluted to 0.48 µg/mL was deposited on freshly cleaved Mica, covered to avoid evaporation and incubated for 40 min at 4-6°C, then rinsed three times by gently exchanging the buffer by pipetting in imaging buffer, before placing in the AFM head for imaging. Scanning was performed at 100 Hz (100kKz PORT rate) covering 512 pixels × 256 lines (for an 800 nm total scan size), corresponding to 2.56 s frame^−1^. Background correction was performed after each frame as described in^2^.

PORT-HS-AFM movies were processed with a custom written plugin in Gwyddion^3^: each image was subjected to plane leveling, initial match line (median of differences row alignment), and background subtraction (flatten base). Images were further subjected to a conservative denoise filter (2 pixels) and saved in the tiff file format.

### Opening and closing lifetime analysis

To automatically detect the lifetimes of open and closed rings, we first manually selected 242 PORT-HS-AFM individual ring sequences from movies exhibiting opening and closing events (see Supplementary Fig. 3a). The movies were imported in IgorPro (WaveMetrics) for further analysis. For each ring, a circular fitting was performed at each frame by computing the sum over a circular profile centered in (*X*_*c*_, *Y*_*C*_) and of radius R for all X_c_ and Y_C_, varying from the center of the image ± 20 pixels to allow for residual imaging drift and ring mobility, as well as for radii varying from R_min_=5 and R_max_=10 pixels, with a step size of 1 pixel. The set of (*X*_*c*_, *Y*_*C*_, R) values maximizing the sum of values above a threshold determined empirically by visual inspection of the quality of the final fit and corresponding to 2.7 nm in height were selected along the circle (see Supplementary Fig. 3b, red ring). The corresponding profiles were plotted over time, and frames for which each point was above the minimal threshold were classified as closed (see Supplementary Fig. 3c). For each automatically detected opening and closing event, a manual validation step was performed thereafter and ring symmetry determined from the number of emanating spokes. Two types of frame sequences were retained for computing lifetimes: i) initially closed rings that opened and then closed again at a later time, without change in symmetry (lifetime open); ii) initially opened rings that closed and then opened again at a later time, without change in symmetry (lifetime closed).

### Oligomer segmentation and classification

Fiji^4^ was utilized to combine images from PORT-HS-AFM movies in a single tiff stack, which was then loaded in the machine-learning software Ilastik^5^. In Ilastik, first brush strokes were drawn to separate each image into 3 channels based on all the available features: 1) background; 2) heads; 3) spokes. The corresponding pixel predictions maps were then utilized for the final segmentation, whereby a simple threshold (T=0.5) in the head channels was applied to segment the different oligomeric entities. This classification process was then utilized to manually label objects and thus train a classifier, also using all available features, for the following 14 classes; oligomers containing 1-10 homodimers, as well of closed rings with 7-10 homodimers. The resulting classification, consisting of the probability of each object to be assigned to each class, was exported as a hd5 file for further processing in IgorPro.

### Kinetics fitting procedure

The output of the classification was used to compute the number of assemblies of each class in every frame as 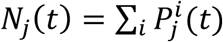, i.e. the probability 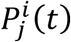 that an object i is in a class j, with i spanning all detected objects in that frame. The error in classification was estimated as 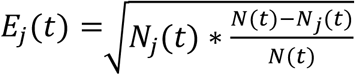, which was derived under the assumption that the variance of a single bin in a multinomial distribution under normal approximation is *Var* (*N*_*j*_(*t*)): = *N*(*t*) * *P*_*j*_(*t*)(1 – *P*_*j*_(*t*)). The resulting kinetics for each PORT-HS-AFM movie were also used to compute the overall concentration of homodimers as the weighted sum of all the classes, i.e. 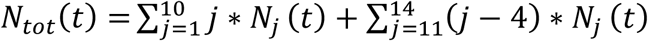. We additionally computed the average polymerization degree as 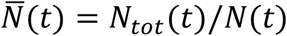 (see Supplementary Fig. 4).

Usually the coagulation-fragmentation equations are considered for closed systems in which the overall mass of the polymers is conserved (see Supplementary Note for a more detailed discussion). However, here we have an open system with a constant increase in polymer concentration over time due to homodimer influx into the chamber and subsequent adsorption to the Mica surface; such an increase needs to be taken into account for the fit. In principle, this increase could have been determined experimentally in each frame. However, to avoid introducing unavoidable noise associated with experimental extrapolations, which would result in non-smooth boundary conditions in the numerical solution of the ordinary differential equations, in the fitting procedure we opted for including the effect of the increase in total concentration over time with an additional terms in the C_1_ concentration (i.e. the concentration of SAS-6 homodimers), accounting for diffusion in solution after injection as well as adsorption and desorption from the surface.

For the concentration in solution we approximated the temporal dependence as:

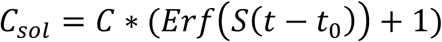

The error function recapitulates the fast increase and subsequent saturation very well. For the surface processes we assume:

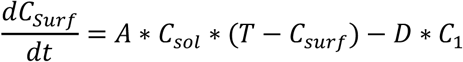

Thus the increase is driven by the concentration in solution C_sol_ and limited by the capacity T of the surface. The surface-adsorbed homodimers then feed into the coagulation-fragmentation equations through C_1_. These equations were then fitted for each PORT-HS-AFM movie to derive the parameters C, S, A, T and D. These parameters were then fixed for the ensuing fitting, which was performed with the global fit package of IgorPro, fitting all the time dependent curve for the 14 species with the same k_on_ and k_off_. It should be noted that the equation for adsorption/desorption does not necessarely reflect the actual adsorption/desorption process and is used here solely as an effective equation fitting to derive accurately the trend of the surface concentration over time.

### Model Comparison

To verify if the proposed coagulation-fragmentation model is adequate for the SAS-6 ring assembly reaction as measured experimentally, we fitted both a simpler model and a more complex one to the measured trajectories. The simplest model that can be considered is one in which only individual homodimers can be added or removed. In this case, which is also known as the Becker-Döring model^6^, the equations simplify to:

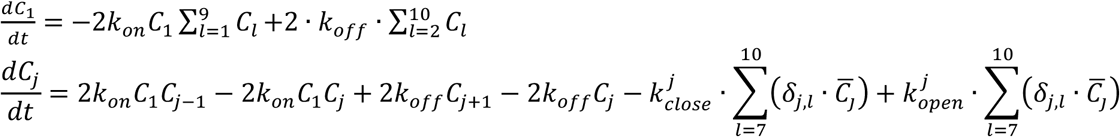

The corresponding fit has a higher reduced Chi-squared than our model (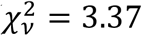 versus 1.01), indicating poorer performance. This had to be expected because visual inspection clearly showed coagulation and fragmentation event of differently sized oligomers. For comparison with a more complex model, instead of the most complex one, which would have required 68 free parameters to allow for all possible interaction constants between all oligomeric species, we opted for an intermediate solution. Considering the fact that homodimers seem to be the species with the largest rotational freedom, we generated a model in which all k_on_ and k_off_ are identical except for those involving encounters with homodimers, which are treated with 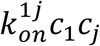, adding 9 more free parameters in total. As expected, the resulting fit yielded a slightly lower reduced Chi-square 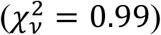. Since the requirements to use Akaike criterion (or Bayesian Information Criterion) are not met in our case given the non-Gaussian distribution of errors, we decided to test if the additional parameters fitted on a single movie were increasing the predictive power of the model on the other movies. Chi-square calculated considering the experimental data and the predicted curves for all movies using the parameters fitted from a single one was not significanly improved by the more complex model 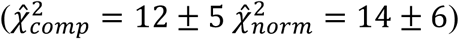, thus validating our model choice.

### Fitting of angle between homodimers

To find the best fitting chain of open SAS-6 oligomeric species, we adopted a three-step approach. First, images of oligomeric species were extracted from the Ilastik output, together with their class prediction. For each particle, a 50×50 pixels^2^ region centered around the geometrical center of the oligomer was cropped and the maximum height normalized. This region was subsequently masked using the head segmentation described in the Oligomer segmentation and classification section, so that only the head domain was retained. In a second step, a custom algorithm written in Python was applied to assign the initial chain position and angles for each cropped oligomer. In brief, the algorithm consists of an optimization performed over the angles between the segments of a continuous line to find the angles that best match the observed image. Initially a chain of N connected vertices was generated at an angle of 40°, where N is the oligomeric state derived from Ilastik. An image was then generated with a line connecting these vertices; this line was then blurred with a gaussian filter to mimic the broadening in the resolution due to imaging. This artificially generated image was then aligned to the original one with the template matching algorithm matchtemplate (OpenCV library), and thus the best relative XY position and corresponding image cross-correlation value identified. The initial angles were then modified by a vector 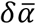 consisting of all inter-segment angles plus the initial angle, thus defining the overall rotation of the entire line. The vector 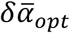 that maximizes the correlation then served as output of the optimization algorithm. The operation was repeated for N-1 and N+1 connected vertex to verify that the maximal correlation corresponded to the predicted class. If that was not the case, the process was iteratively repeated, starting from the N* maximizing the correlation. Given the stochastic nature of minimum finding algorithms in general, the line fitting procedure was further refined in a third step by an exhaustive search algorithm written in IgorPro, starting from the solution found in the previous step, which was subjected to iterative small angle modifications spanning all angles ±10° (every 1°). At each iteration with an angle modification, the two successive angles were re-calculated to solve the corresponding four-bar linkage problem^7^. The angle over which the image profile was maximized was retained and kept as the starting condition for the following angle until angles were stable.

### Spoke mobility analysis from PORT-HS-AFM

To analyze spoke mobility following PORT-HS-AFM imaging, we first manually selected 10 9-fold rings that remained closed for >20 frames. For each frame of each ring, we then generated a circular profile, fitting the head domains with the algorithm described in the Ring Opening and Closing Analysis section. From those positions, we generated a second profile with a radius larger by 10 nm (see Supplementary Fig. 9a). These outer profiles were then used to generate kymographs, which were manually divided into 9 regions corresponding to the 9 spokes (see Supplementary Fig. 9b). The location of each spoke was then tracked across the kymograph by finding the maximum height within the relevant region in each frame. A time window of 13 frames was then picked randomly from each kymograph to avoid averaging spoke behavior that might have changed over longer times. Within this time window, discrete spokes positions in each frame were used to compute the average displacement over time from the average spoke position. The histogram of the average square displacement for all spokes in the 10 selected rings is shown in Supplementary Fig. 9c. The average displacements for each spoke were then classified based on a manually selected threshold into immobile or mobile on the surface (see Supplementary Fig. 9e). To verify the stability of this analysis, we computed at varying threshold the number of spokes classified as immobile, as well as the number of equally oriented spokes (see Supplementary Fig. 9d, green line). We then confronted this computation with the number of equally oriented spokes calculated from a random distribution of immobile and mobile spokes, using varying probability for each spoke to be mobile. We found that the experimentally measured distribution differs significantly from a random one over a sizeable range (see Supplementary Fig. 9f).

### Molecular dynamics

We modeled a solution structure of the *Chlamydomonas* SAS-6[6HR] nonamer based on the PDB structures 3Q0Y and 3Q0X^1^, which contain respectively two head domains connected by their N-terminal interface (3Q0Y) and a homodimer including 6 heptad repeats of the coiled-coil interface (3Q0X). Since 3Q0Y has a higher resolution (2.1 Å compared to 3.02 Å of 3Q0X), we built a homodimer homology model from the 3Q0Y head domain and the 3Q0X tail domain by aligning the two structures using UCSF Chimera (version 1.12^8^) and Swiss Model in user template mode^9^. To generate a nonameric assembly, we then aligned a head domain of 3Q0Y to a head domain of the new hybrid homodimer model and used this as an N-terminal interface template to align a second homodimer structure. This procedure was repeated until a complete nonamer was generated, which exhibited a spiral structure. The martinize.py script (version 2.6,^10^) was used to obtain the structure in coarse-grain representation.

To build a flat nonameric SAS-6 ring, we used Gromacs tools version 2018^11,12^ to align one coarse-grained homodimer with the xy-plane. We copied this homodimer and rotated each subsequent copy by 40° within this plane until nine homodimers were arranged in a flat circle, with the N-terminal head interfaces of two neighboring homodimers close to each other in the center and the 6 heptad repeats of the coiled-coil pointing outwards. We then utilized an elastic network to close the ring. Elastic networks are commonly utilized in coarse-grain simulations to compensate for the loss of detailed non-covalent interactions^13^. We obtained the elastic network by coarse-graining a SAS-6[6HR] dimer using martinize.py, with 0.5 nm and 0.9 nm for the lower and upper cut-offs, respectively, and replicating it to have the same network on all nine interfaces. The same elastic network was also imposed on the spiral structure for stabilization. To pull the nine homodimers arranged in the above manner into a closed ring, we ran a 1 ns simulation with 5% (25 kJ mol^-1^ nm^-2^) of the original force constant on the elastic network. The complete simulation conditions are listed below. To get SAS-6[6HR] oligomers smaller than a nonamer from both the flat ring and the spiral shaped structure, the required number of homodimers was deleted from the respective nonameric structures to obtain the desired size. For simulations including a generic surface, we only used the structures derived from the flat ring; the surface was generated using VMD (version 1.9.3^14^) Nanotube Builder with carbon beads placed with graphene geometry at a distance of 0.282 nm.

All coarse-grain simulations were performed with Gromacs version 2018.5 along with the Martini force field (version 2.6)^10–12^. We based the surface parametrization on non-bonded C1 beads, but decreased the interaction strength between protein and surface beads by two levels in the Martini interactions table, and increased the water-surface interactions from a repulsive to an intermediate interactions level. This parametrization has a lower hydrophobicity, and allows diffusion on the time scale of the MD simulations, resulting in a surface with medium-level adhesion strength of the protein, i.e. a generic flat template for SAS-6[6HR]. For simulations of SAS-6[6HR] in solution, we placed the protein structures in a cubic box with 3 nm minimal distance to the nearest box edge. For simulations on a surface, we chose the x- and y-dimensions to fit the generic surface (46×46 nm). The size in the z-direction was 20 nm. All systems were solvated with Martini water, spiked with 10% anti-freeze water and neutralized by the addition of 150 mM NaCl. We energy minimized the systems using the steepest descent algorithm with a step size of 0.01 nm until energy convergence and a maximal force below 100 kJ mol^-1^ nm^-1^. Surface beads were frozen during minimization and position restrained by a harmonic potential with a 1000 kJ mol^-1^ nm^-2^ force constant during all other equilibration and production simulation steps.

We equilibrated the minimized structures for 1 ns in the NVT ensemble, followed by 200 ps in the NPT ensemble with a 2 fs integration time step, as well as 1 ns in the NPT ensemble with a 10 fs time step and a 1000 kJ mol^-1^ nm^-2^ force constant on protein position restraints. To allow further relaxation of the structures, we ran two 30 ns simulations with a 30 fs time step and decreased force constants for position restraints on protein beads of 200 and 100 kJ mol^-1^ nm^-2^, respectively. The production simulations each continued for 1.5 µs with a 30 fs integration time step without position restraints on protein beads. Periodic boundary conditions were applied in all simulations. We kept the temperature at 320 K using the velocity-rescaling thermostat with a coupling constant of 1.0 ps. Isotropic pressure coupling was ensured by the Parrinello-Rahman barostat with a compressibility of 3.0×10^−4^ bar^-1^ and a coupling constant of 12.0 ps. We treated long-range interactions using a reaction-field, assuming a dielectric constant of infinity beyond a cutoff of 1.1 nm and a relative dielectric constant of 15. Van-der-Waals interactions were cut off after 1.1 nm. The Verlet neighbor list was updated every 20 integration steps, and we set the Verlet-buffer-tolerance to 0.005 kJ mol^-1^ ps^-1^.

We prepared the SAS-6[6HR] dimer structure in all-atom resolution based on the crystal structures as described above and placed it in a dodecahedron simulation box with a minimal distance from the nearest protein atom to the box edge of 2 nm. We solvated with TIP3P water and neutralized by the addition of 150 mM NaCl. We energy minimized the system using the steepest descent minimization with a step size of 0.01 nm until energy convergence and a maximal force below 1000 kJ mol^-1^ nm^-1^. For all simulations at all-atom resolution we used the Amber99sb*ildnp force field^15^ with a time step of 2 fs for the leap frog integrator. We equilibrated for 100 ps in the NVT ensemble followed by 1 ns in the NPT ensemble. A constant temperature of 300 K and pressure of 1 bar was ensured respectively by the velocity-rescaling thermostat with a coupling constant of 0.1 ps and Berendsen barostat with a coupling constant of 2 ps and a compressibility of 4.5×10^−5^ bar^-1^. We cut off all short range interactions after a distance of 1 nm and treated the long range electrostatic interactions using the Particle-Mesh-Ewald method. The Verlet neighbor list was updated every 20 integration steps and periodic boundary conditions were in place. All bonds involving hydrogen atoms were restrained using the LINCS algorithm. Positions restraints with a 1000 kJ mol^-1^ nm^-2^ force constant, which we used to restrain all protein heavy atoms during equilibration, were released for the subsequent production simulation runs. These continued for 200 ns each with the same conditions as applied to the equilibration steps except that we used the Parrinello-Rahman barostat with a coupling constant of 5 ps for isotropic pressure coupling.

For the analysis of angles between neighboring homodimers, vector 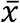 describes the orientation of the head domain and is defined as the vector connecting the center of mass of both head domains, while vector 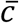 describes the orientation of the 6 heptad repeats and connects residue F153, which marks the beginning of the coiled-coil between the two head domains of a homodimer, and residue Q391 close to the C-terminal end of the 6 heptad repeats (Supplementary Fig. 8). Vector positions were extracted using Gromacs (version 2018) tools and subsequent geometric analysis was done using custom python 3 scripts. The in-plane angles describe hinge movement of two homodimers a and b relative to each other. The inplane angle *α* is defined as the angle between 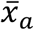 and 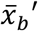, with 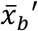 being the projection of 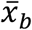 into the plane defined by 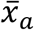 and 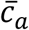. The torsion angle *η* is defined as the angle between 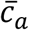 and the surface.

To analyze whether the distribution of spokes is random or correlated, we iterated over the neighbors of each spoke and calculated the probability for the neighbors to be in the same orientation, i.e. bound or not bound, based on the overall percentage of bound or unbound spokes. We compared the resulting number of spokes expected to have the same orientation as their neighbor given a random distribution to the number of spokes that were actually observed to have the same orientation.

Spoke mobility was calculated by a root mean square fluctuation (RMSF):

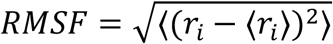

with *r*_*i*_ being the position of residue Q391 within the xy-plane or along the z-axis. The spokes were classified as bound or not bound based on a position cut-off in z-direction.

## Acknowledgments

We thank E.A. Lemke, P. Guichard, F. Schneider and A. Bezler for critical reading of the manuscript. This work was supported by the European Research Council (ERC), though AdG 835322, CENGIN to P.G., as well as StG 307338, NaMic and CoG 773091, InCell to G.E.F. Further support came from the Swiss National Science Foundation through the European Union’s Seventh Framework Programme FP7/2007-2011 under grant 200021_182562 agreement 286146 to G.E.F. N.B. was supported by the EPFL Fellows postdoctoral fellowship program funded by the European Union’s Horizon 2020 Framework Program for Research and Innovation (Grant agreement 665667, MSCA-COFUND). F.G., S.d.B. and U.S.S. acknowledge funding through the Deutsche Forschungsgemeinschaft (DFG, German Research Foundation) under Germany’s Excellence Strategy – 2082/1 – 390761711. F.G. and S.d.B. acknowledge funding by the Klaus Tschira Foundation, the state of Baden-Württemberg through bwHPC, as well as the DFG through grant INST 35/1134-1 FUGG. S.d.B. thanks the Carl Zeiss Foundation for financial support.

