## Supplementary Materials for "Surface-catalyzed SAS-6 self-assembly directs centriole formation through kinetic and structural mechanisms"

**Supplementary Note**

### Coagulation-fragmentation equations for SAS-6 ring assembly

#### General remarks

Association and dissociation processes are ubiquitous in nature and can be described mathematically by the coagulation-fragmentation equations[1,2]. If association and dissociation only occur in steps of monomer addition or removal, these equations are known as the Becker-Döring equations[3,4]. Because it was originally conceived for processes that are dominated by association rather than dissociation and that can grow to very large sizes (e.g. growth of ice crystals, river deltas or planets), the mathematical treatment of these equations often focuses on the limit of large systems.

In the context of growth of supramolecular assemblies in biological systems, the dissociation process tends to be of the same nature as the association process; in particular, the binding and unbinding of proteins involves the association and dissociation of the same contacts in the binding interface. Moreover, system size tends to be finite and limited by some maximum. Predictions of the coagulation-fragmentation equations have been shown in the case of SAS-6 ring formation to agree with the results from Brownian dynamics computer simulation of patchy particles forming five- or nine-rings[5,6]. Because the transport process underlying association in such a patchy particle model is purely diffusive, it was possible to demonstrate that perfect agreement with the coagulation-fragmentation equations can be achieved by appropriately incorporating the diffusion process into the association rates of the coagulation-fragmentation equations, thereby combining spatial and kinetic modeling. In fact, the diffusion constants also entered in the dissociation rates due to the condition of detailed balance. This earlier work now allows us to study the cases of diffusion-limited and reaction-limited regimes. In each case, one can identify the time scales for the growth dynamics and the steady states. In addition, one can study the role of absorbing boundaries, e.g. a stabilizing effect of ring closure that arises when a sufficiently large number of proteins have assembled.

Here we show that in the reaction-limited regime, the coagulation-fragmentation equations simplify to a form that effectively depends only on one unknown parameter, the ratio of bond association and dissociation rates, which can be extracted from experimental data and converted into an effective equilibrium constant. We also discuss the deviations that occur due to transport processes.

#### Coagulation-fragmentation equations

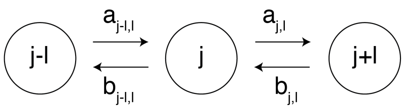


**Fig. SN1: Reaction scheme for the coagulation-fragmentation equations.**

*Association rates $a_{j,l}$ and dissociation rates $b_{j,l}$ are labeled by indices that denote which fragment sizes are involved. Note that in principle the rates could be very different dependent on the fragment size, because the reactions of differently size fragments can involve very different transport and reaction processes. The special case $l = 1$ corresponds to a monomer addition and removal scheme, in which case the coagulation-fragmentation equations are called Becker-Döring equations.*

We first introduce the coagulation-fragmentation equations in their general form and then specify them for the purpose of the SAS-6 assembly process. We follow the nomenclature from[2]. The concentration of a fragment of size j is denoted by $c_j$. As shown in Fig. SN1, we have to consider two association and two dissociation processes that can change the concentration of a cluster of size j (with $j \geq 2$). Using the law of mass action and summing over all possible sizes of the reaction partners, we obtain the following system of ordinary differential equations:

**Supplementary Equation (1)**

$$\dot{c}_j(t) = \frac{1}{2}\sum_{l=1}^{j-1} a_{j-l,l}c_{j-l}c_l - \sum_{l=1}^{N-j} a_{j,l}c_jc_l - \frac{1}{2}\sum_{l=1}^{j-1} b_{j-l,l}c_j + \sum_{l=1}^{N-j} b_{j,l}c_{j+l}$$

The signs in front of the four different terms denote if the concentration $c_j$ increases or decreases by the respective process. Usually the coagulation-fragmentation equations are studied in the limit $N \to \infty$, but here we consider a maximal cluster size $N$.

The first term describes the coagulation of a $(j - l)$-cluster with a $l$-cluster into a $j$-cluster. The factor of $1/2$ reflects that the sum includes two terms that are equivalent; e.g. both a $(1,4)$- and a $(4,1)$-encounter can give a 5-cluster, but they represent the same process. Alternatively, one could have left out the factor of $1/2$ and restrict the summation to one of the two pairs[5,6]. The second term describes the coagulation of a $j$-cluster with a $l$-cluster into a $j + l$ cluster. In both cases, the association rate $a_{j,l}$ describes the probability with which a cluster of size $j$ encounters and reacts with a cluster of size $l$. Obviously, we have $a_{j,l} = a_{l,j}$. Because of the product of the two concentrations due to mass action, these equations are non-linear. In three dimensions, the physical dimension of the association rates $a_{j,l}$ is $1/Ms$ or $m^3/s$. In two dimensions, it would be $m^2/s$. These dimensions reflect the fact that coagulation is bimolecular and depends on the concentration of a partner. In particular, this implies a transport process that brings a partner to close proximity before binding can occur.

The third term in Supplementary Equation (1) describes how a $j$-cluster decays into a $(j-l)$ −cluster and a $l$-cluster. Again, we have a factor of $1/2$ to avoid overcounting. The fourth term describes the decay of a $(j+l)$ cluster into a $j$-cluster and a $l$-cluster, thus increasing the number of $j$-clusters. For the dissociation rate, we also have $b_{j,l} = b_{l,j}$ irrespective of the exact mechanism of fragmentation. In both three and two dimensions, the physical dimension of the dissociation rates $b_{j,l}$ is $1/s$. This reflects the fact that fragmentation is a unimolecular reaction, starts from a situation of close encounter and does not depend on concentration of any partner.

We note that the case of two equally sized clusters requires extra care[5]. For association between $n$ equally sized clusters, there are $n(n-1)/2$ possible pairings. Thus, in the thermodynamic limit of large system size, when the law of mass action becomes valid and when n-1 ≈ n, this requires a factor of $1/2$, which is already contained in the notation of Supplementary Equation (1).

For $j = 1$ and $j = N$, obviously two of the four processes are not possible and the corresponding terms will be missing. For these special cases, sometime the rates are redefined with a factor of 2 (e.g. $a_{1,1} = 2a_1$ for the Becker-Döring equations), to obtain a more symmetric definition of the system of equations in terms of fluxes[7].

One important consistency check of the coagulation-fragmentation equations from Supplementary Equation (1) is that because they are formulated for a closed system here, they should obey mass conservation based on the symmetry of the matrices $a_{j,l}$ and $b_{j,l}$:

$$\sum_{j=1}^{N} j\,\dot{c}_j(t) = 0$$

The simplest test example would be N = 3, the formation of a trimer. Then only four reactions are possible: 1+1 →2; 2+1 → 3; 3 → 2+1; 2 →1 + 1. The system of ordinary differential equations following from Supplementary Equation (1) reads:

$$\dot{c}_1(t) = -a_{1,1}c_1^2 - a_{1,2}c_1c_2 + b_{1,1}c_2 + b_{1,2}c_3$$

$$\dot{c}_2(t) = \frac{1}{2}a_{1,1}c_1^2 - a_{1,2}c_1c_2 - \frac{1}{2}b_{1,1}c_2 + b_{1,2}c_3$$

$$\dot{c}_3(t) = a_{1,2}c_1c_2 - b_{1,2}c_3$$

One sees that these equations are non-linear because the monomer concentration $c_1$ varies in time. Importantly, one also can verify that mass conservation is obeyed, $\sum_{j=1}^{3} j\,\dot{c}_j(t) = 0$, and identify the different terms that balance each other after multiplication with the monomer number $j$ per cluster. For the case N=10 studied here, there are many more terms, thus we do not document them here explicitly. Using computer algebra, one can verify that they obey mass conservation as expected.

**Effect of transport**

In the coagulation-fragmentation equations from Supplementary Equation (1), association is described by an association rate $a_{j,l}$ that in general depends on the cluster sizes $j$ and $l$. These rates are determined by two aspects: first a transport mechanism has to bring the two clusters in close spatial proximity, and, second, a reaction can then take place to generate the product. Here we consider diffusion to be the relevant transport process. To determine the relative importance of diffusion versus reaction, we write this sequence of events in the following scheme[5] (for simplicity, we drop the indices and consider $j$ and $l$ to be fixed):

$$A + B \underset{b}{\overset{a}{\rightleftharpoons}} C \;, A + B \underset{k_{Diff,b}}{\overset{k_{Diff,a}}{\rightleftharpoons}} AB \underset{k_d}{\overset{k_a}{\rightleftharpoons}} C$$

Here A and B are two oligomers coagulating to a new oligomer C. Conversely, the oligomer C can fragment into the two oligomers A and B. The overall association rate $a$ and the overall dissociation rate $b$ are the ones that should enter the coagulation-fragmentation equations. $k_{Diff,a}$ and $k_{Diff,b}$ denote the diffusive association and dissociation rates, respectively, which have dimensions of $m^3/s$ ($m^2/s$ in two dimensions) and $1/s$ (independent of dimension), respectively. AB is the encounter complex separating the transport step from the reaction step. $k_a$ and $k_d$ are the association and dissociation rates, respectively, for the reaction from encounter of the complex to the final product; both have dimension $1/s$. These four rates can be identified in computer simulations of the patchy particle model.

For sufficiently good statistics, we will assume the encounter complex $AB$ to be in steady state. From this we obtain the effective association and dissociation rates[7,8]:

**Supplementary Equation (2)**

$$a = \frac{k_{Diff,a} k_a}{k_a + k_{Diff,b}}, \; b = \frac{k_{Diff,b} k_d}{k_a + k_{Diff,b}}$$

This in turn allows us to identify the dissociation constant, which is also the concentration at half-occupancy:

**Supplementary Equation (3)**

$$K_d = \frac{b}{a} = \frac{k_{Diff,b} k_d}{k_{Diff,a} k_a} = \frac{k_d}{V^* k_a}$$

Here we have identified the reactive volume $V^* = k_{Diff,a}/k_{Diff,b}$ as the physical factor which is required to convert the non-spatial rate $k_a$ for bond formation into an association constant that reflects the concentration dependance. For simple cases, the reactive volume can be calculated analytically; for more complex cases, it has to be determined through computer simulations. For proteins like SAS-6, the typical value is set by the size of the contact region and therefore has a typical value below $nm^3$. For the five- and nine-ring configurations, the reactive volumes have been tabulated for all possible cluster combinations $(j, l)$[5,6].

Supplementary Equation (2) allows us to identify the two regimes of reaction and diffusion control. In the reaction-limited regime ($k_a \ll k_{Diff,b}$), we get $a = V^* k_a$ and $b = k_d$. We further specify the situation of patchy particles that have two opposing patches that lead to oligomerization, as is the case for the SAS-6 homodimers. Then the reaction volume is twice the reactive volume of one patch, $V^* = 2V_p^*$, and the microscopic association rate $k_a$ is the rate of bond formation when two patches encouter each other in space. Thus $a = 2V_p^* k_a$. The microscopic dissociation rate can be identified with twice the internal rate of bond dissociation, $b = 2k_d$, because there are two ways to cut a $(j + l)$-cluster into a $j$-cluster and a $l$-cluster. For equally sized clusters, there is only one way to cut, but then the double number of clusters is produced, thus a factor of 2 is justified in this case as well. Now the coagulation-fragmentation equations form Supplementary Equation (1) simplify to:

**Supplementary Equation (4)**

$$\dot{c}_j(t) = V_p^* k_a \sum_{l=1}^{j-1} c_{j-l}\, c_l - 2V_p^* k_a c_j \sum_{l=1}^{N-j} c_l - k_d(j-1)c_j + 2k_d \sum_{l=1}^{N-j} c_{j+l}$$

This is exactly the form of the coagulation-fragmentation equations used in the main text, if one identifies $k_{on} = V_p^* k_a$ and $k_{off} = k_d$. Because for the patchy particles we also have investigated previously the role of diffusive transport, we now can address the effect of making the assumption of a reaction-limited process.

#### Formation of a SAS-6 nine ring without closure

We next discuss the assembly of a nine-fold ring (N=9), which is the case of the SAS-6 system (without ring closure or considering other ring symmetries). In contrast to the experimental situation, but in agreement with the usual situation studied with the coagulation-fragmentation equations, we consider that we start with a certain amount of monomers, which then are continuously used up. Using molecular information and affinity measurements from the literature, this system has been parametrized and compared to patchy particle simulations before [6]. Because assembly in solution at the centrosomal concentration of c = 5 μM (corresponding to e.g. 125 particles in a box of linear size 346 nm) was found in computer simulations to be very slow (time scale of days), a parametrization with enhanced reactivity but with the same known dissociation constant has been developed, which here we use as a reference case. In this parametrization, we have $V^* = 0.0585\ nm^3$, $k_a = 4$ / ns and $k_d = 1.06\ 10^{-6}$ / ns. Using Supplementary Equation (3), this leads to $K_d = 7.5$ μM for the patchy particles, which in turn corresponds to the experimentally measured $K_d = 60$ μM for single N-term domains (factor 8 due to combinatorics of patches). The diffusive on-rate has been simulated to be $k_{Diff,a} = 3\ 10^5$ / M s for the SAS-6 homodimers, which is three orders of magnitude smaller than the Smoluchowski rate for isotropic proteins and suggests that assembly of SAS-6 in solution is diffusion-limited in this case[9]. Indeed, the corresponding diffusive off-rate of our parametrization is $k_{Diff,b} = k_{Diff,a} / V^* = 8.55\ 10^{-3}$ / ns, which is much smaller than $k_a$. To explore the reaction-controlled regime, one has to lower both $k_a$ and $k_d$ by three orders of magnitude, so that $k_a$ becomes smaller than $k_{Diff,b}$ and $K_d = k_d / (V^*\ k_a)$ stays constant at the experimentally known value.

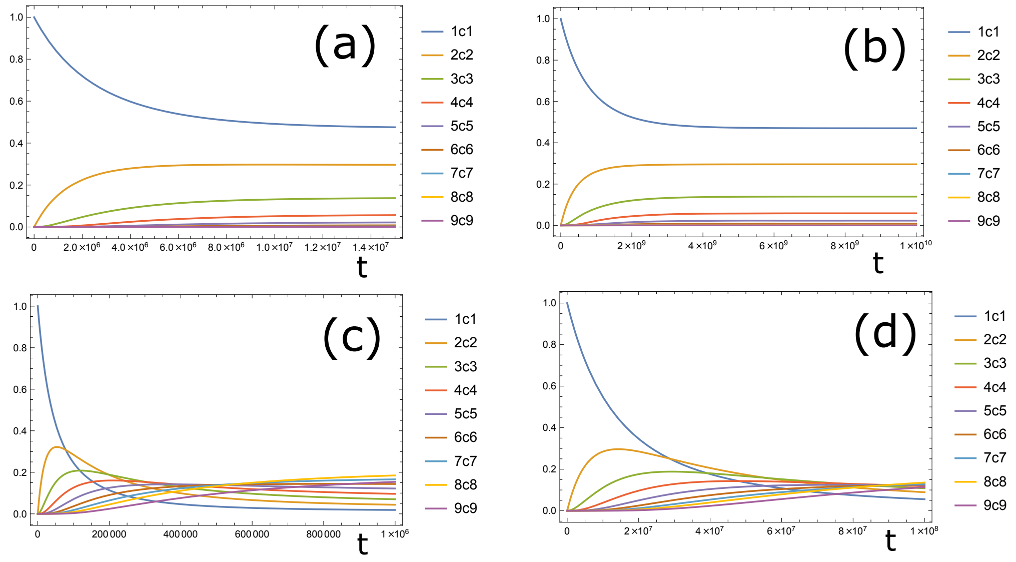


**Figure SN2: Formation of a nine-fold SAS-6 ring in solution using the coagulation-fragmentation equations taking into account diffusive transport and with the parametrization of enhanced reactivity from[6].**

*Time t measured in ns, concentrations $c_j$ normalized by $c_1$. (a) Reaction rates $k_a$ = 4 / ns and $k_d$ = 1.06 $10^{-6}$ / ns, concentration c = 5 μM, reactive volume for homodimers V* = 0.0585 $nm^3$, dissociation constant $K_d$ = 7.5 μM, diffusive association rate $k_{Diff,a}$ = 3 $10^5$ / Ms. At this centrosomal concentration, no rings and only small oligomers are formed in solution. The system is diffusion-limited. (b) When both $k_a$ and $k_d$ are reduced by three orders of magnitude, the system becomes reaction-limited and much slower (note the different scale on the time axis), but the dynamics is qualitatively very similar. (c) For strongly increased concentration c=500 μM, rings are formed even in solution. Otherwise parameters as in (a). (d) Again, going to reaction-control by decreasing both $k_a$ and $k_d$ by three orders of magnitude like in (b) does not change the dynamics qualitatively, except that the dynamics becomes much slower.*

In Figure SN2 we plot the time evolution that one gets for these parameter values, using the exact values for the reactive volumes and diffusive rates obtained in calibration simulations for the patchy particles and tabulated in [5]. Figure SN2(a) corresponds to Figure S4 of [6] and demonstrates that in solution no rings are formed even at centrosomal concentration (time measured in ns). Because this situation is strongly diffusion-controlled, in Figure SN2(b) we show that reaction-control does not really change this picture, except making the dynamics much slower. Formation of rings can be obtained only at high concentration, as shown in Figure SN2(c), where he have increased it by a factor of 100 compared to (a). In (d), we show that also in this case, reaction-control qualitatively gives very similar results (except making assembly much slower again). We conclude that the assembly dynamics of SAS-6 is mainly determined by concentration and less by whether it is diffusion- or reaction-controlled.

**Ring closure**

We finally add the important aspect that after reaching the upper size limit $N$, the ring can undergo a further step of closure. This step is characterized by a rate $k_a^{intra}$ that does not depend on transport, but should be determined by the energy gained during ring closure. Because experimentally we have observed rings with sizes ranging between 7 and 10, we now set $N = 10$ and allow for an additional (closed) state of all clusters with $7 \leq j \leq 10$. This leads to the final form of our equations:

$$
\begin{aligned}
\dot{c}_j(t) &= k_{on} \sum_{l=1}^{j-1} c_{j-l}\, c_l - 2k_{on} c_j \sum_{l=1}^{N-j} c_l + 2k_{off} \sum_{l=1}^{N-j} c_{j+l} - k_{off}(j-1)c_j \\
&\quad - k_{close}^{j} \sum_{l=7}^{10} \delta_{jl}\, c_j + k_{open}^{j} \sum_{l=7}^{10} \delta_{jl}\, \overline{c}_j \\
\dot{\overline{c}}_j(t) &= -k_{open}^{j} \overline{c}_j + k_{close}^{j} c_j
\end{aligned}
$$

The first equation is valid for $2 \leq j \leq 9$. For $j = 1$ and $j = N$, two of these terms are missing. The second equation is valid for $7 \leq j \leq 10$.

**Supplementary Note References**

**Supplementary Fig. 1: CrSAS-6 constructs and purification**

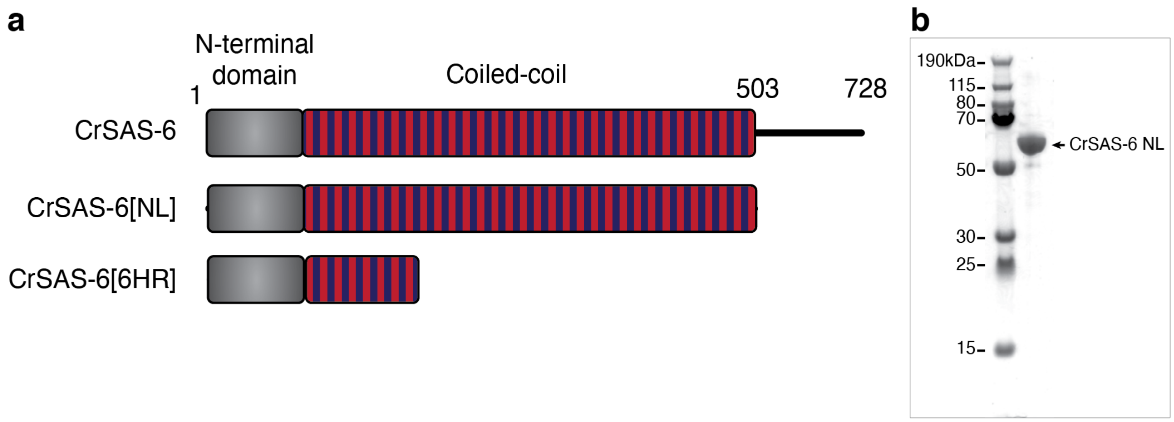


**a**, Schematic of the full length 728 amino acid long CrSAS-6 protein (top), the CrSAS-6[NL] protein used in the PORT-HS-AFM experiments (middle), and the CrSAS-6[6HR] protein used for the MD simulations (bottom).
**b**, Comassie staining of SDS-PAGE with recombinantly expressed and purified Cr-SAS-[NL], with an indication of molecular weight markers.

**Supplementary Fig. 2: SAS-6 rings of 7- to 10-fold symmetries**

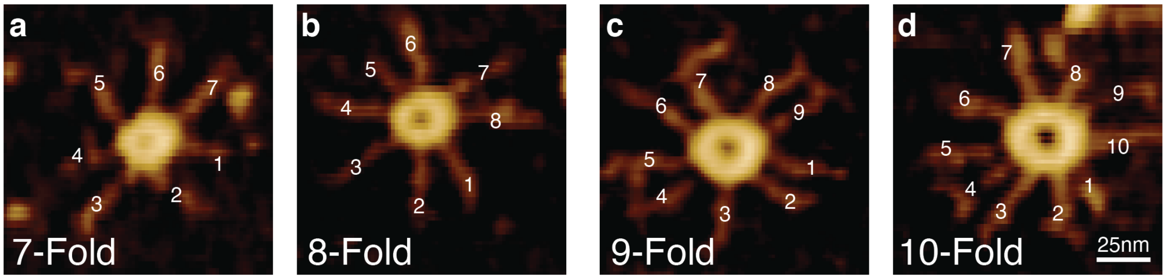


**a-d,** Representative images of SAS-6 rings of indicated symmetries (determined by the number of spokes) observed by PORT-HS-AFM. Note that ring diameter increases with higher fold-symmetries.

**Supplementary Fig. 3: Ring opening and closing analysis.**

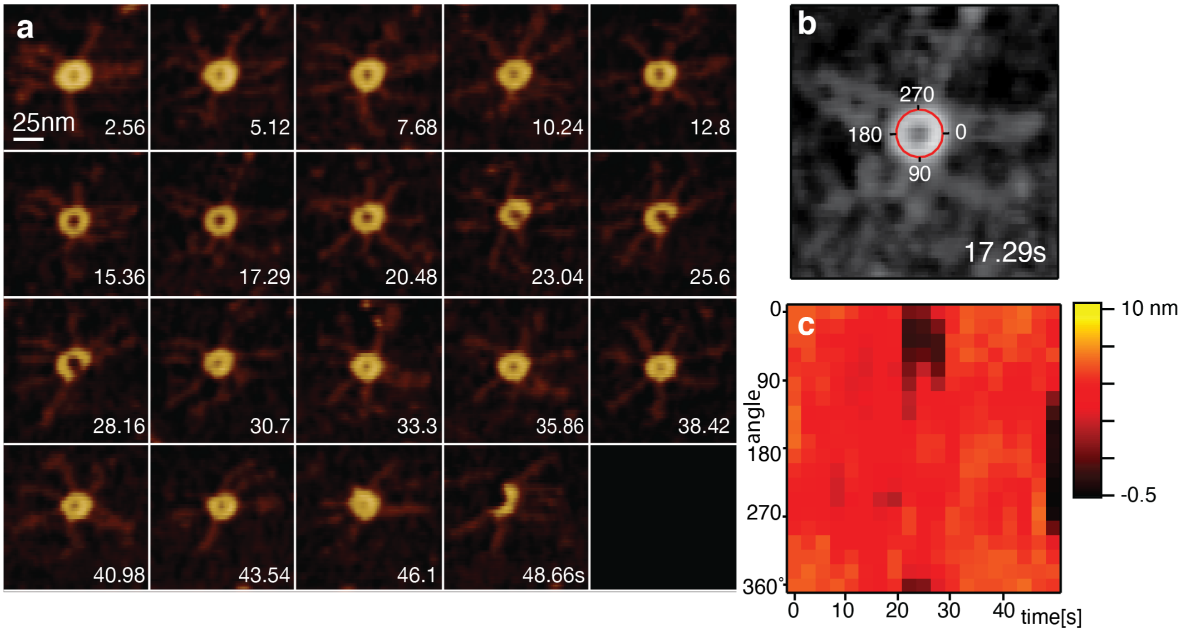


**a**, Imaging of SAS-6 ring with PORT-HS-AFM, from full ring to complete disruption (time is indicated in seconds; 2.56 s frame rate). Open rings are observed at 23.04, 25.6 and 28.16.
**b**, Circular profile (red line) fitted to the ring formed by the head domains.
**c**, Corresponding heights along the circular profile over time, which allows detection of ring opening events as regions with lower height (in this case between frames 23.04 and 28.16), and to then manually identify opening and closing transitions.

**Supplementary Fig. 4: Segmentation and classification pipeline.**

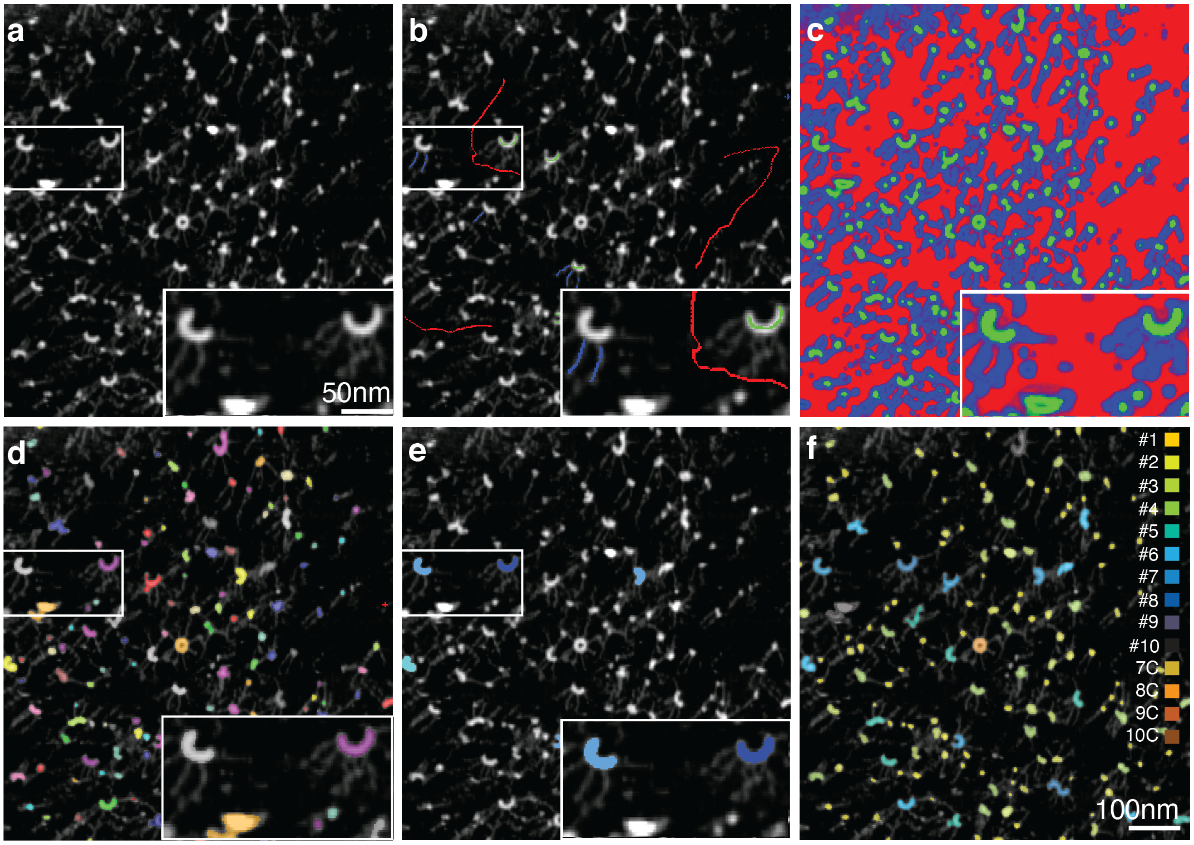


**a**, Individual frame from a SAS-6 PORT-HS-AFM experiment. High magnification inset is shown on the bottom right for this and other panels in this figure.
**b**, Different brushes drawn to identify background (red), head domains (green) and spokes (blue).
**c**, Pixel prediction map representing background (red), head domains (green) and spokes (blue).
**d**, Head domains within SAS-6 oligomers identified after thresholding. Colors are assigned randomly to the distinct oligomeric assemblies.
**e**, Manual training of selected object into one of 14-classes (1-10 open oligomers, plus 7- to 10-fold rings). Some oligomers with clearly identifiable spoke numbers are assigned to a specific class at this step (e.g. colored oligomers in the frame)
**f**, Final classification result with probability of belonging to a certain class for each detected object. The class with the highest probability is displayed in the corresponding color.

**Supplementary Fig. 5: SAS-6 assembly kinetics.**

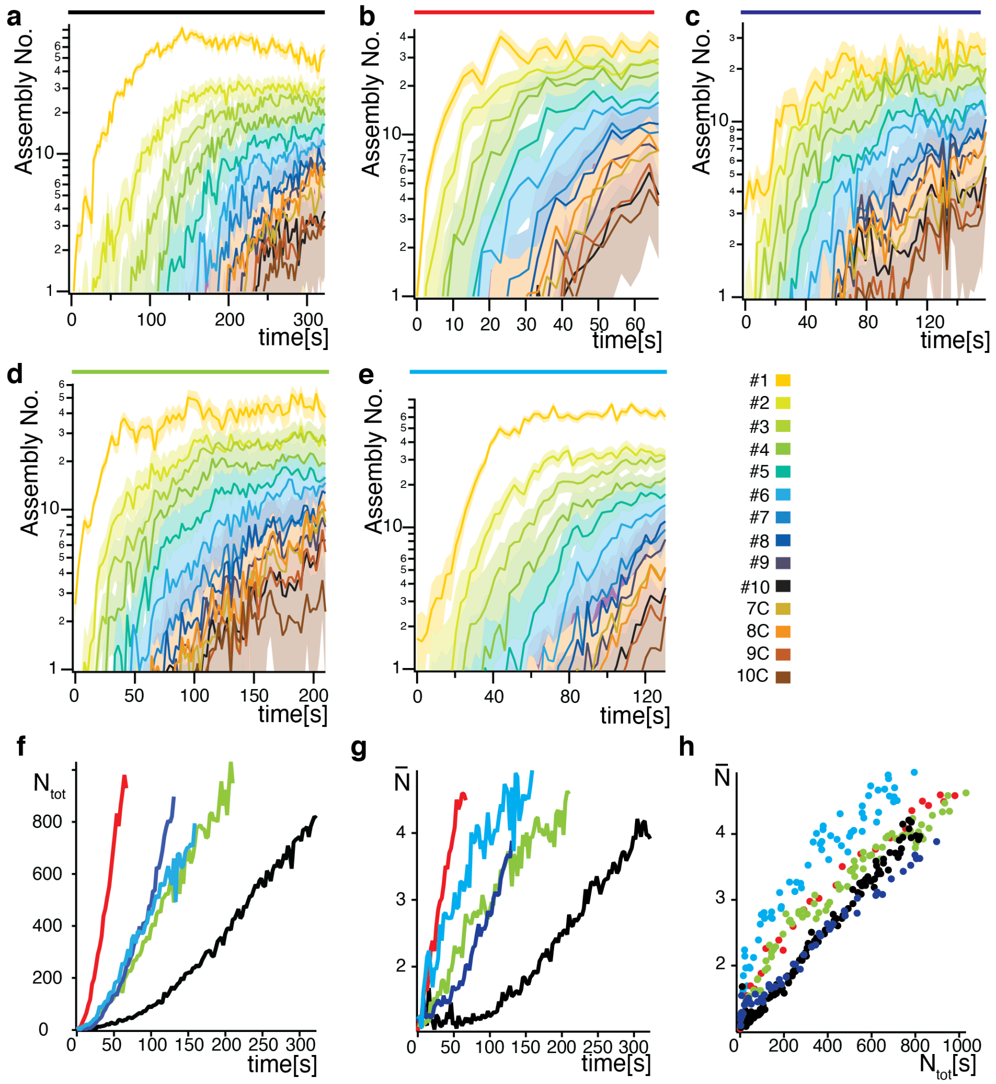


**a-e**, SAS-6 assembly kinetics resulting from the segmentation output of the five PORT-HS-AFM experiments analyzed in this study; A is the same as in Fig. 3b. Colors above the plot correspond to lines shown in F-H.

**f**, Total number of homodimers independently of their oligomeric state ($N_{tot}$) on the Mica surface over time for the five PORT-HS-AFM experiments, each represented with a different color. The diversity in the traces reflects variable rates of homodimers adsorption on the surface, likely arising from variability in protein injection in the measurement chamber.

**g**, Average oligomerization degree $\bar{N}$ over time for the five PORT-HS-AFM experiments. The variation in adsorption kinetic is reflected in the variable polymerization degree over time.

**h**, Average oligomerization degree ($\bar{N}$) as a function of total number of homodimers on the Mica surface. Note similar slopes, reflecting analogous oligomerization kinetics.

**Supplementary Fig. 6: Overall effective surface concentration and model comparisons**

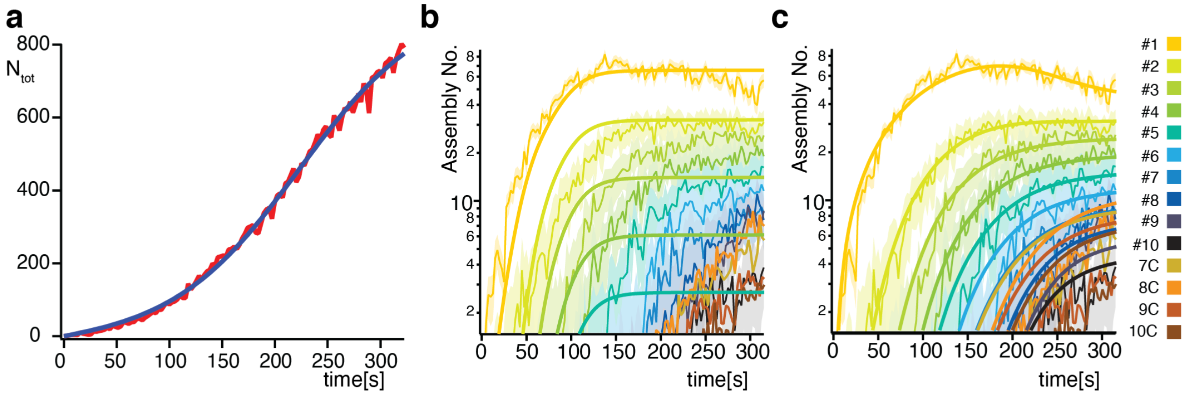


**a**, Total number of adsorbed homodimers on the Mica surface in the PORT-HS-AFM experiments independently of their oligomeric state ($N_{tot}$) as a function of time (red), with corresponding fit (blue), for the experiment shown in Fig. 3b and Supplementary Fig. 5a.
**b**, Fit of the kinetics for the experiment shown in Fig. 3b and Supplementary Fig. 5a using the simpler Becker-Döring model, in which only association and dissociation events of individual homodimers are considered.
**c**, Fit of the kinetics for the experiment shown in Supplementary Fig. 3b and Supplementary Fig. 5a using a model with more free parameters, in which all $k_{on}$ and $k_{off}$ are identical except for those involving encounters with homodimers (see Materials and Methods).

**Supplementary Fig. 7: Determination of angles between SAS-6 homodimers**

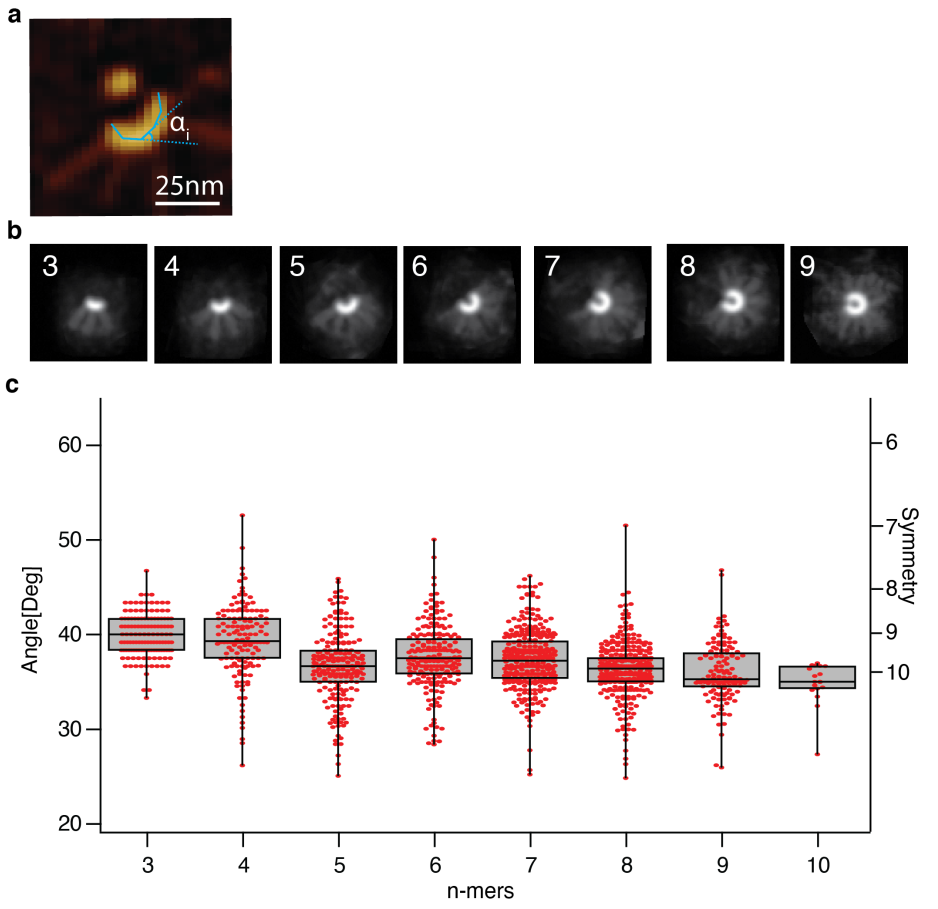


**a**, Scheme of poly-line fit performed on PORT-HS-AFM data, with $\alpha_i$ being the fitted angle.
**b**, Average images of each re-aligned open oligomeric species, based on the poly-line fit in the PORT-HS-AFM experiments (note that the low number of 10-mers was not sufficient to generate a meaningful average configuration).
**c**, Box plot of angles and predicted symmetries retrieved from fitting.

**Supplementary Fig. 8: Additional features of molecular dynamics simulations**

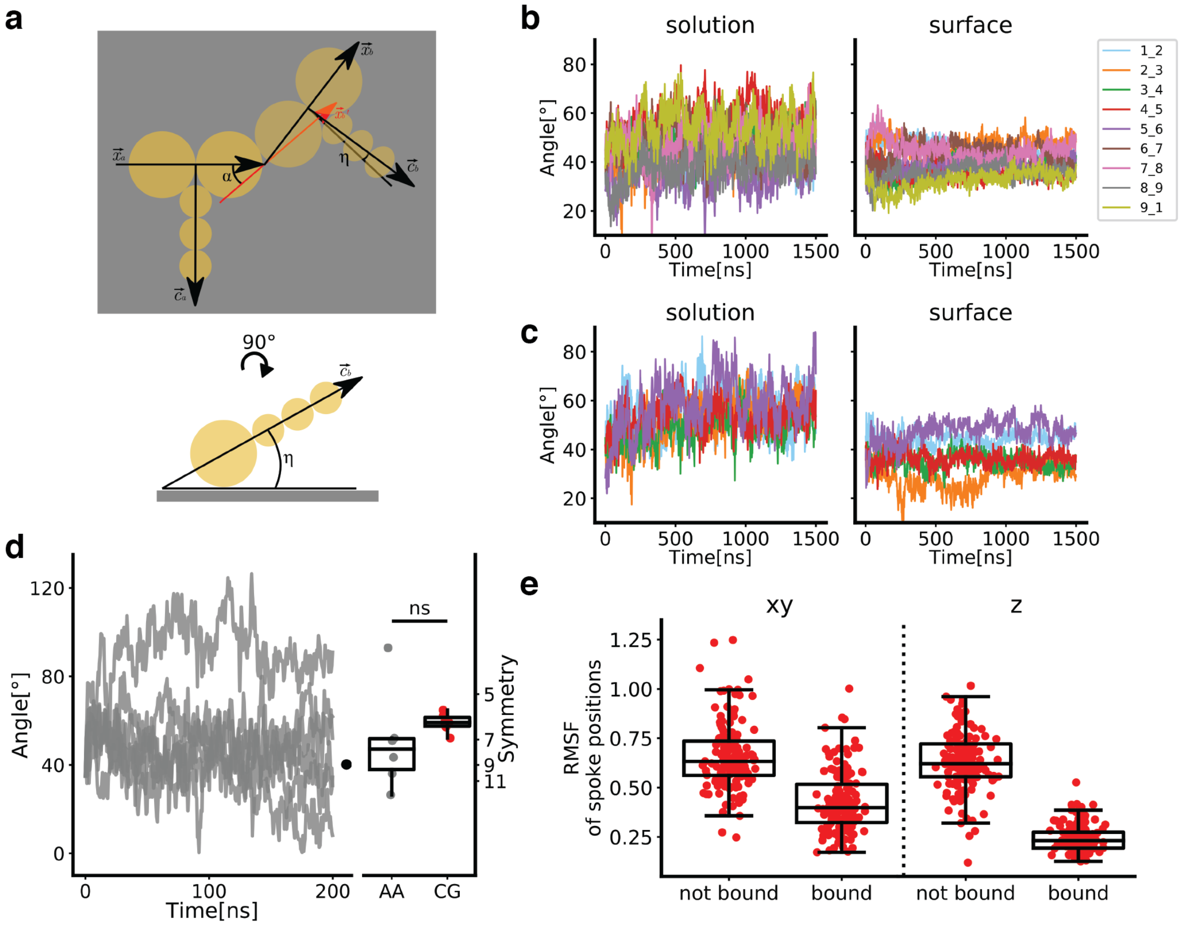


**a**, Schematic of measured angles between SAS-6 homodimers; α: in-plane angle; η: torsion angle with respect to the surface.

**b**, **c**, Individual time traces of the in-plane angles for nonamer (**b**) and hexamer (**c**), on a surface and in solution.

**d**, Time traces (grey plots) and averages (grey scatters) of in-plane angles in all-atom (AA) simulations of SAS-6[6HR] dimers, revealing the large range of angles explored by SAS-6 and therefore the need perform extensive sampling. Average angles measured in coarse-grain (CG, red scatters) simulations of SAS-6[6HR] dimers, as well as in a simulation of an N-terminal head domain dimer (black scatter) from [18], which both fall within this range. ns: not significant, p=0.38

**e**, In-plane (xy) and off-plane (z) mobilities of spokes bound or not bound to the surface.

**Supplementary Fig. 9: Spoke mobility analysis**

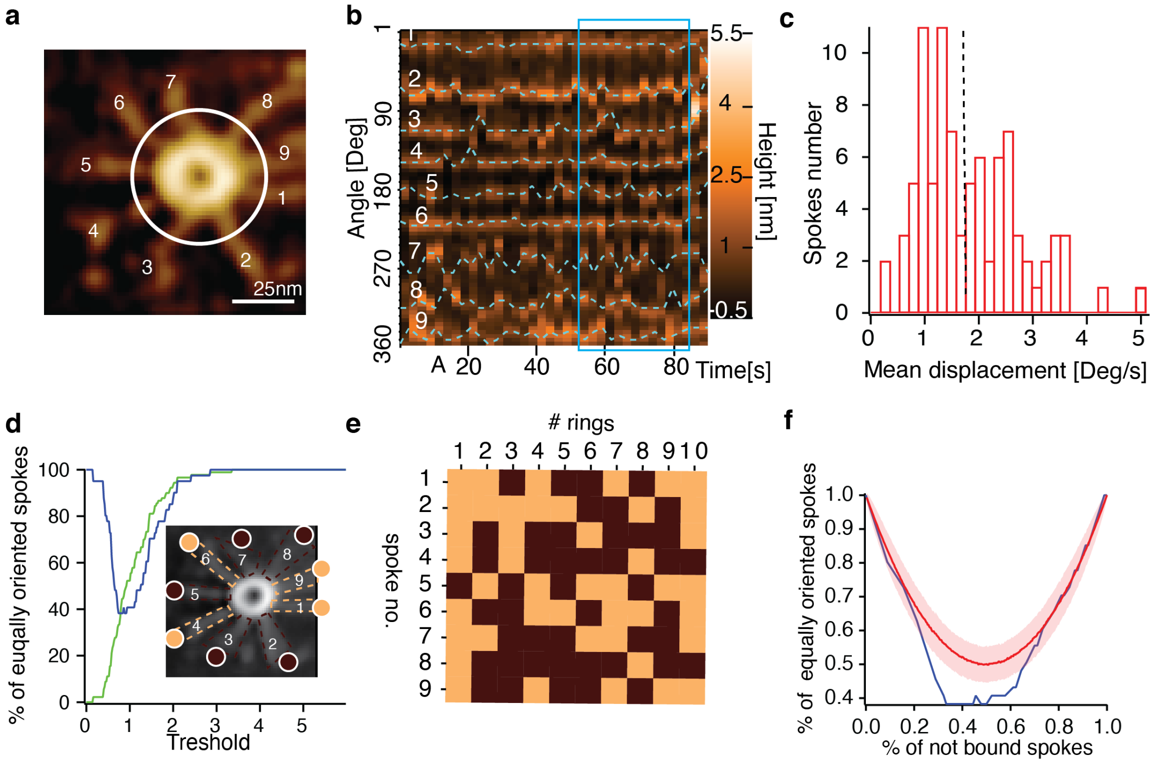


**a**, PORT-HS-AFM frame illustrating circle utilized to construct kymograph shown in B.
**b**, Kymograph of spoke angle position and corresponding tracks over time (blue dashed lines). Mean displacement was derived from a randomly selected 13-frame block (blue rectangle).
**c**, Distribution of spoke mobility from 10 closed 9-mer rings monitored by PORT-HS-AFM; dash line indicates threshold used to discriminate between mobile and immobile spokes.
**d**, Experimentally determined fraction of spokes immobile (green line) at increasing thresholds of spoke mobility set to discriminate immobile versus mobile spokes, with corresponding fraction of spokes oriented in the same manner, either immobile or mobile (blue line). Inset shows an average image from a 13-frame block with spokes classified as immobile (beige dashed lines and disks) or mobile (dark brown dashed lines and disks) at the selected threshold.
**e**, Matrix representation of bound or not bound spokes for 10 closed 9-mer monitored by PORT-HS-AFM. The distribution is not compatible with a random distribution ($p<0.022$).
**f**, Fraction of spokes with similar mobility as a function of the fraction of mobile spokes expected from a random distribution determined from a Monte-Carlo simulation (red line, with SD), and from the experimentally determined distribution (blue line), where the variable fraction of mobile spokes (X axis) arises from varying the mobility threshold (see panel D). This analysis shows that the distribution is statistically different (out of the 2-sigma threshold) from the random one in a significant interval (% of not bound spokes from 0.35-0.59).

**Supplementary Table 1:**

| | 7 closed | 8 closed | 9 closed | 10 closed |
|---|---|---|---|---|
| **7 closed** | NA | <0.001 | 0.004 | NS |
| **8 closed** | <0.001 | NA | NS | NS |
| **9 closed** | 0.004 | NS | NA | NS |
| **10 closed** | NS | NS | NS | NA |
| | **7 open** | **8 open** | **9 open** | **10 open** |
| **7 open** | NA | 0.02 | 0.004 | NS |
| **8 open** | 0.02 | NA | 0.014 | NS |
| **9 open** | 0.004 | 0.04 | NA | NS |
| **10 open** | NA | NA | NS | NA |

Significance difference (p-values) for mean open and closed lifetimes between symmetries as indicated (tested with F-ratio test on the mean values).

**Supplementary Movie 1: SAS-6 oligomerization and ring assembly**
HS-PORT-AFM movie showing progressive adsorption of recombinantly expressed SAS-6 homodimers on the Mica surface, their progressive oligomerization and then ring formation. The color scale represents height and is the same as that reported in Fig. 1.

**Supplementary Movie 2: SAS-6 ring opening and closing**
Example HS-PORT-AFM movie of a SAS-6 ring preformed on Mica opening, closing and eventual disassembly. The color scale represents height and is the same as that reported in Fig. 1.

**Supplementary Movie 3: Classification result from machine learning software Ilastik**
Visualization of machine learning class prediction for the different assembly state of oligomers present in Movie S1. The overlaid colors represent the assigned oligomerization state following the color code depicted in Fig. 3b.
